# Cryo-EM structure of Pol κ−DNA−PCNA holoenzyme and implications for polymerase switching in DNA lesion bypass

**DOI:** 10.1101/2020.07.10.196956

**Authors:** Claudia Lancey, Muhammad Tehseen, Masateru Takahashi, Mohamed A. Sobhy, Timothy J. Ragan, Ramon Crehuet, Samir M. Hamdan, Alfredo De Biasio

**Affiliations:** Leicester Institute of Structural & Chemical Biology and Department of Molecular & Cell Biology, University of Leicester, Lancaster Rd, Leicester LE1 7HB, UK; Division of Biological and Environmental Sciences and Engineering, King Abdullah University of Science and Technology, Thuwal 23955, Saudi Arabia; CSIC-Institute for Advanced Chemistry of Catalonia (IQAC) C/ Jordi Girona 18-26, 08034 Barcelona, Spain

## Abstract

Replacement of the stalled replicative polymerase (Pol δ) at a DNA lesion by the error-prone DNA polymerase κ (Pol κ) restarts synthesis past the lesion to prevent genome instability. The switching from Pol δ to Pol κ is mediated by the processivity clamp PCNA but the structural basis of this mechanism is unknown. We determined the Cryo-EM structures of human Pol κ–DNA–PCNA complex and of a stalled Pol δ–DNA–PCNA complex at 3.9 and 4.7 Å resolution, respectively. In Pol κ complex, the C-terminus of the PAD domain docks the catalytic core to one PCNA protomer in an angled orientation, bending the DNA exiting Pol κ active site through PCNA. In Pol δ complex, the DNA is disengaged from the active site but is retained by the thumb domain. We present a model for polymerase switching facilitated by Pol κ recruitment to PCNA and Pol κ conformational sampling to seize the DNA from stalled Pol δ assisted by PCNA tilting.

---

Cells are continously subjected to DNA damage caused by environmental mutagens and reactive metabolites, which threaten the stability of the cell genome^1,2^. At a DNA lesion, the cell faces a choice between stalling DNA replication or employing a more error-prone replication system that tolerates the damage before it can be repaired. Translesion DNA synthesis (TLS) is the process that allows cells to overcome the deleterious effects of replication stalling and genomic instability caused by DNA damage^3–6^. While being of the utmost importance for cell survival, TLS is also intrinsically mutagenic and is implicated in human cancer^7–9^. Eukaryotic TLS involves canonical high-fidelity as well as specialised error-prone TLS polymerases (*e.g*., Y-family Pol η, Pol ι, Pol κ and Rev1) which can synthesise DNA past a lesion due to their active site being able to accommodate a damaged template^3,10,11^. Recent work showed that, in yeast, the lagging strand Pol δ is responsible for lesion bypass on both leading- and lagging strands. The leading strand Pol ε dissociates from the DNA upon encountering a lesion and travels with the CMG helicase to support uncoupled leading strand synthesis. Pol δ is recruited to the lesion site until a TLS reaction bypasses the lesion and synthesis by Pol δ resumes before it is replaced by Pol ε and the replisome is restored^12^. Both Pol δ and TLS polymerases form complexes with the homotrimeric sliding clamp PCNA, which encircles duplex DNA and tethers these polymerases to the template, enhancing their catalytic rate and processivity^13–15^. At the lesion site, Pol δ stalls and PCNA is mono-ubiquitylated at K164 by Rad6–Rad18 ubiquitin ligase complex^16–18^; a TLS polymerase then binds to the resident PCNA and replicates the damaged DNA^19^. PCNA ubiquitylation facilitates the recruitment and retention of TLS polymerases to damage sites *in vivo*^20–26^ and in a fully reconstituted yeast replisome^12^. The structural basis of the interaction of TLS polymerases with PCNA, and the mechanism of exchange with the high-fidelity polymerase during TLS are poorly understood.

Eukaryotic Y-family polymerases display significant functional divergence, making them highly specialized for the bypass of specific lesions^27^. Pol κ can bypass several types of damage mainly at the N^2^ position of guanine in an error-free manner^28^, and efficiently extend mispaired termini with lower misincorporation frequency than undamaged templates^29^. While Pol κ lacks the ability to insert nucleotides opposite the 3’T of a *cis-syn* thymine dimer, it can extend past a dG inserted opposite the 3’T of the dimer by another DNA polymerase (e.g., Pol η)^29^. In addition, Pol κ can generate single-base frame-shifts through template–primer misalignments^27,30^. Recently, it has been shown that Pol κ is able to exchange with Pol δ that is stalled at repetitive common fragile sites^31^.

Pol κ differs from Pol η and ι in that its orthologs exist in bacteria and archaea, including DinB (Pol IV) in *Escherichia coli* and Dbh and Dpo4 in *Sufolobus solfataricus*^28^. Nonetheless, it shares a similar domain architecture with Pol η and ι consisting of an N-terminal catalytic domain (comprising a palm, fingers, thumb, and PAD) and a long C-terminal domain containing two PCNA-interacting motifs (PIP-boxes), one Rev-1 interacting motif (RIR) and two Ubiquitin Binding Zinc Fingers (UBZs), and predicted to be largely unstructured (Figure 1a). An extension of ∼75 amino acids at the N-terminus (N-Clasp), which is functionally important, is a unique feature of Pol κ^32^. The structure of the catalytic domain of human Pol κ has been solved in the apo form and in complex with DNA^32,33^. The apo and DNA-bound structures of Pol κ display a large difference in the orientation of the PAD relative to the thumb domain. In the apo enzyme, the PAD is positioned under the palm domain in two alternate positions, while in the DNA-bound form it is docked in the major groove; for the most divergent position, a movement requiring a 50 Å shift and a 143° rotation^32^ (Figure 1b). Conformational freedom of the PAD in the apo form has also been observed in Dpo4^34^ and, to a minor extent, in Pol η^35,36^, and seems to be a general feature of Y-family polymerases. DNA binding to Pol κ also results in the folding of the N-Clasp into two helices (αN1 and αN2) encircling the primer template junction^32^.

**Figure 1.**
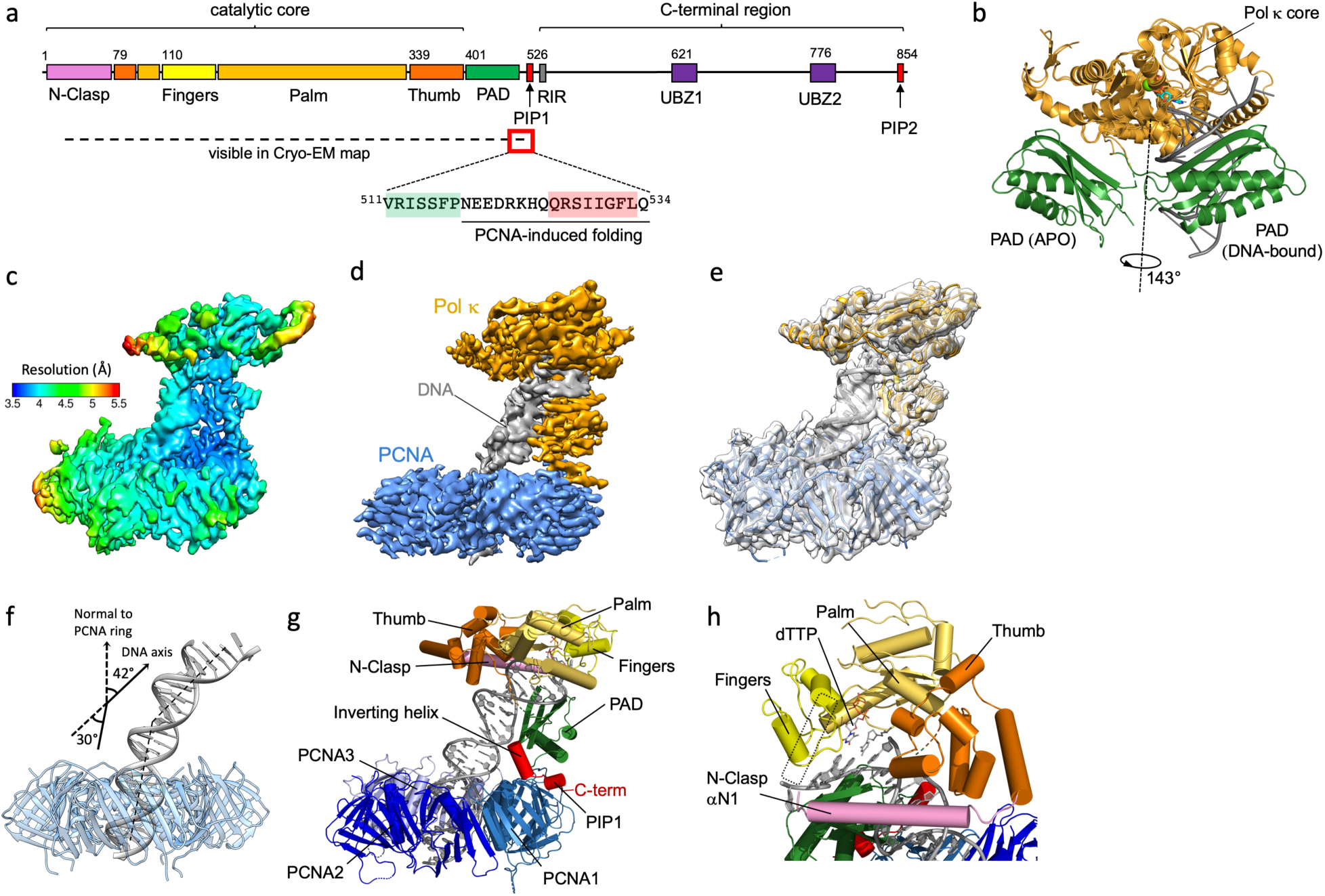
(a) Cryo-EM structure of Pol κ–PCNA–DNA complex. **a)** Domain organization of human Pol κ and amino acid sequence of the PCNA-interacting region; PIP: PCNA interacting motif; RIR: Rev-1 interacting motif; UBZ: Ubiquitin binding zinc-finger. **b)** X-ray structures of apo-(PDB ID 1T94)^33^ and DNA-bound Pol κ (PDB ID 2OH2)^32^ overlaid on the core domain (orange ribbon). The PAD domain (green ribbon) in the apo structure is rotated 143° relative to the PAD in the DNA-bound structure. **c**) Cryo-EM map of Pol κ holoenzyme color-coded by local resolution. **d)** Cryo-EM density map of Pol κ holoenzyme complex colored by components (Pol κ in orange, PCNA in skyblue and DNA in grey). **e)** Pol κ holoenzyme structure fitted into the Cryo-EM map. **f)** DNA bending in Pol κ holoenzyme model. PCNA is shown as a transparent blue ribbon, DNA as a grey ribbon. Pol κ core and PAD domain were removed for clarity. **g)** Structure of Pol κ holoenzyme colored by domain. **h)** Close-up of Pol κ catalytic core colored by domain. The dotted rectangle corresponds to the position of N-Clasp αN2 in the X-ray structure of Pol κ ternary complex (PDB ID 2OH2)^32^, which is invisible in the Cryo-EM map.

The interaction of Pol κ with PCNA is principally mediated by the two PIP-box motifs at the C-terminus of Pol κ, each responsible for a specific function^20^. While both PIP-boxes promote PCNA ubiquitylation, only the internal PIP-box stimulates DNA synthesis by Pol κ. In addition, PCNA ubiquitylated at K164 slightly enhances Pol κ synthetic activity compared to unmodified PCNA^20^, suggesting a weak but significant interaction between the UBZ zinc fingers of Pol κ and the ubiquitin moieties of ubiquitylated PCNA. Accordingly, PCNA ubiquitylation was shown to be important to recruit Pol κ to stalled replication forks^37–39^.

Mammalian Pol δ consists of a catalytic subunit (p125) and three regulatory subunits (p50, p66 and p12), all required for optimal holoenzyme activity^40^. We have recently determined the Cryo-EM structure of human Pol δ–PCNA–DNA complex captured in the act of synthesis, showing that the p125 subunit binds to one PCNA protomer in an open conformation and the regulatory subunits are positioned laterally^41^. This arrangement allows PCNA to thread the P/T DNA exiting the catalytic cleft while exposing its unoccupied monomers to recruit other proteins, as demonstrated for flap endonuclease 1 (FEN1), the enzyme responsible for cleavage of the 5’ flap of Okazaki fragments^41^. Importantly, the Pol δ–DNA–PCNA complex exists in different conformations with increased loosening of the interactions with PCNA and increased tilting of PCNA, which may further expose the clamp to accommodate bulky partners^41^.

In this work, we present the 3.9 Å-resolution Cryo-EM structure of human full-length Pol κ bound to PCNA, a P/T DNA substrate, and an incoming nucleotide, which we refer to as Pol κ holoenzyme. Pol κ is docked to one PCNA protomer through the internal PIP-box adjacent to the PAD domain. The region C-terminal to the PIP-box and containing the UBZ zinc fingers, is instead invisible in the Cryo-EM map. Pol κ holoenzyme architecture shows an unusual arrangement, where the catalytic domain and the DNA exiting Pol κ core are sharply angled relative to the PCNA ring. Our MD simulations predict that, in the absence of DNA, Pol κ bound to PCNA is highly flexible, suggesting that binding to DNA is required for the assembly of the rigid and active holoenzyme. In addition, we present the Cryo-EM structure of a stalled human Pol δ–PCNA–DNA complex, determined at 4.7 Å resolution, in which the P/T DNA is outside the catalytic site but remains attached to the thumb domain of the polymerase. In light of these structures, and the previously determined structures of alternate Pol δ holoenzyme conformers^41^, we propose a mechanism for the handoff of a lesioned DNA substrate between Pol δ and Pol κ, resulting in lesion bypass and restart of replication.

## Results

### Cryo-EM structure of Pol κ holoenzyme

We reconstituted Pol κ holoenzyme by mixing purified recombinant Pol κ, PCNA, a (25/38) P/T DNA substrate containing a dideoxy chain terminator in the primer strand, and dTTP as the incoming nucleotide; the purified Pol κ is active, stimulated by PCNA and able to bypass an abasic lesion (Supplementary Figure 1). The complex was then separated by micro-size exclusion chromatography (Supplementary Figure 2), vitrified and imaged by Cryo-EM (Supplementary Figure 3). We obtained a reconstruction of the complex at a global resolution of 3.9 Å (Figure 1c-e; Supplementary Figure 4, Supplementary Table 1). The structure has approximate dimensions of 127.7 Å × 88.0 Å × 89.1 Å, and displays the catalytic core of Pol κ sitting on top of the front face of PCNA in a remarkably angled orientation, with the axis of DNA in the catalytic cleft tilted by ∼42° relative to the normal of the PCNA ring plane (Figure 1f). The duplex DNA emerging from the catalytic core threads through the PCNA ring hole and bends by ∼30° to avoid clashing with the ring inner rim (Figure 1f). The long region C-terminal to the PAD domain (residues 535-870, Figure 1a) is invisible in the map, suggesting it is disordered. Fitting of the X-ray structure of Pol κ catalytic domain bound to DNA (PDB ID 2OH2)^32^ into the Cryo-EM map shows excellent correlation for Pol κ core, PAD and P/T DNA in the active site, while bending of dsDNA in Pol κ holoenzyme results in poor fitting of the bases below the PAD domain (Supplementary Figure 5). The map quality allowed us to build an atomic model of the full holoenzyme (Figure 1g). Residues 21-45 in the N-Clasp are invisible in the Cryo-EM map (Supplementary Figure 5), suggesting that helix αN1 in the N-Clasp is flexible even in the presence of DNA (Figure 1h). This agrees with the high average B-factor of residues 21-44 in the X-ray structure (107.1 Å^2^) compared to the overall value (69.5 Å^2^), and with the fact that αN1 in the N-Clasp engages in marginal interactions with DNA^32^.

The Cryo-EM map resolution at the Pol κ–PCNA interface (∼3.6 Å; Figure 1c) was sufficient for de-novo model building of this region (Figures 1g and 2a-b). Pol κ interacts with one of the three PCNA protomers mainly through the C-terminal region of the PAD spanning residues 517-534, which is disordered in the absence of PCNA^32^ and becomes structured in the complex. Specifically, residues 518-525 fold into a 2-turn α-helix (“Inverting helix”, Figure 1g) which reverses the chain direction and inserts the PIP-box (^526^QRSIIGFL) between the loop connecting helix αQ and β11 of the PAD, the hydrophobic cleft on the front face of PCNA, and the PCNA C-terminus (Figure 2a-b). The PIP-box acquires the canonical 3.10 helix conformation and docks to the PCNA groove via a 3-fork plug made of side chains of Ile529, Phe532 and Leu533, while Gln526 binds in the so called “Q-pocket” (Figure 2a-b). While deviating from the strict PIP-box consensus sequence (*Qxxhxxaa*, where *h* is a hydrophobic, *a* is an aromatic, and *x* is any residue), Pol κ PIP-box interacts through the prototypical molecular surface observed in other PCNA-interacting partners^42^. Additional interactions further stabilize the structure: two main-chain hydrogen bonds between Pol κ residues Gln526 and Arg527 and Ile255 Pro253 in the C-terminus of PCNA, and two hydrogen bonds between His44 on a PCNA loop adjacent to the hydrophobic cleft and residues Ser528 and Ile529 within Pol κ PIP-box (Figure 2b). Thus, the folding and concomitant insertion of the PAD C-terminus between the PAD and PCNA creates a composite interface burying a total of 890 Å^2^ (567 Å^2^ and 323 Å^2^ of the PCNA and PAD surface areas, respectively), and bends Pol κ core over the PCNA bound protomer, causing the bending of DNA threading the PCNA pore. Interestingly, both Pol κ and Pol δ interact with only one PCNA protomer via a PIP-box interaction involving the C-terminus of the catalytic domain, but the polymerase chains approach the PCNA binding groove from opposite directions, and are connected N-terminally to distinct domains (Figure 2c).

**Figure 2.**
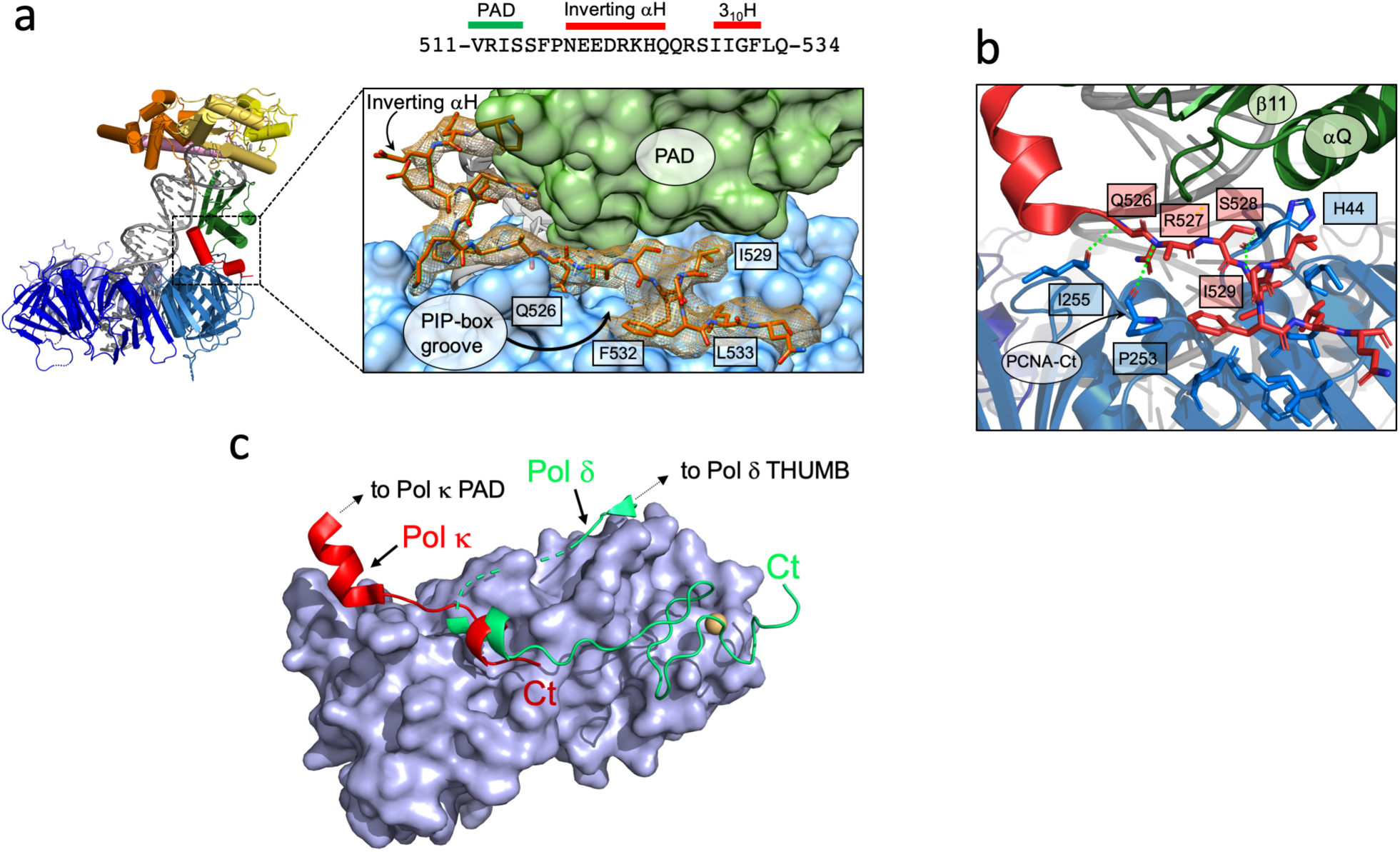
Cryo-EM density and model of Pol κ region interacting with PCNA. **a)** Map region around the PAD C-terminus. PAD C-terminus is shown as a red stick. PCNA and Pol κ PAD are shown as cyan and green surfaces, respectively. Key residues in Pol κ PIP-box are labelled. The amino acid sequence of Pol κ interacting region is shown above the figure, and secondary structure elements are labeled **b)** Inter-molecular interactions. Pol κ PIP-box and PCNA interacting residues are shown as red and blue sticks, respectively. Hydrogen bonds are shown as green dotted lines. Residues forming the canonical PCNA hydrophobic cleft are shown but not labeled for clarity. Pol κ PAD and PCNA are shown as ribbons and colored by domain. **c)** Pol κ and Pol δ binding to PCNA. The region of Pol κ and Pol δ interacting with PCNA are shown as red and green ribbons, respectively. Interacting PCNA protomer is shown as a light blue surface.

### Interaction with DNA in Pol κ holoenzyme

In the active site of Pol κ, Watson-Crick base pairing is observed between the terminal A in the primer strand and the incoming dTTP (Figure 3a-b). The triphosphate of dTTP inserts between the palm and fingers domain and its position is locked by hydrogen bonding with Tyr111, Arg144 and Lys328, three conserved residues among Y-family polymerases (Figure 3c). The side chains of the residues responsible for catalysis (Asp107, Asp198 and Glu199) protrude between the triphosphate portion of dTTP and the phosphate group of the terminal templating base (Figure 3c). The map resolution prevented to discriminate the two metal cations (Ca^2+^) which are normally coordinated in the active site of replicative polymerases. Density at the 5’ end of the template strand is compatible with two bound nucleotides (Figure 3b). The nucleobase of A at position −1 is packed against the Phe49 in αN2 of the N-Clasp, while the nucleobase of T at position – 2 is in close proximity to Ser47, which is the last visible residue of the N-clasp (Figure 3c). This reinforces the notion that the interaction with DNA is important to stabilize the N-Clasp, resulting in the full encirclement of P/T within Pol κ core^32^. Most of the interactions stabilizing the Pol κ–DNA complex involve the PAD domain, and are analogous to those reported in the X-ray structure of Pol κ ternary complex^32^.

**Figure 3.**
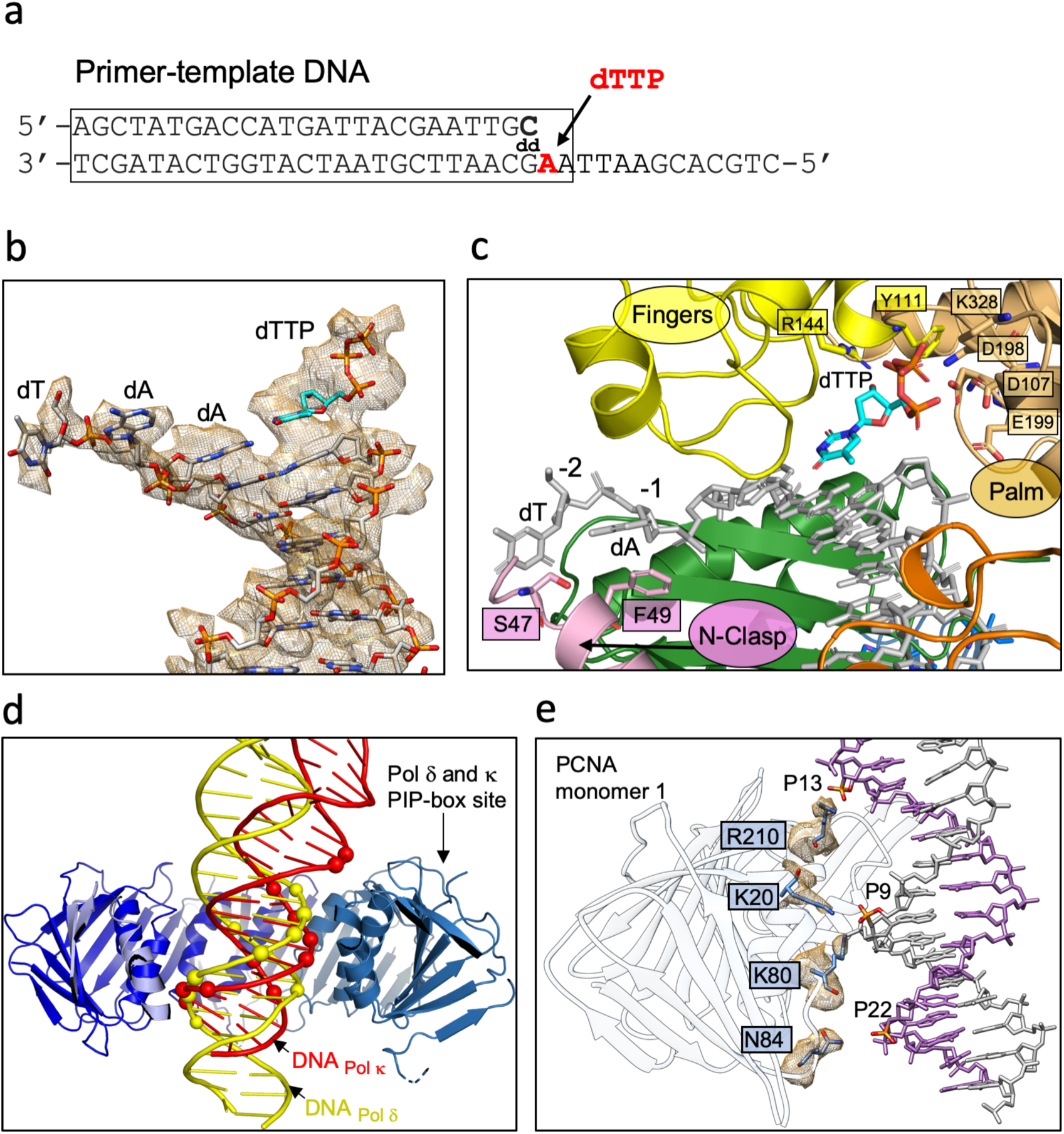
Cryo-EM density and model of Pol κ holoenzyme regions interacting with DNA. **a)** Sequence of the DNA P/T substrate. The region of the substrate that was modelled is boxed. **b)** Map region around P/T DNA in Pol κ active site. **c)** Model region of Pol κ active site. Pol κ is shown as a ribbon colored by domains, residues interacting with DNA are shown as sticks. DNA is shown as grey sticks. **d)** Side-view of the Cryo-EM structures of Pol δ (PDB ID 6TNY)^41^ and Pol κ holoenzymes aligned on PCNA. PCNA subunits are shown in different shades of blue and the subunit in the foreground is removed for clarity. DNA molecules in Pol δ and κ structures are shown as yellow and red ribbons, respectively. DNA phosphates within a coulombic interaction distance (<6 Å) from PCNA polar residues are shown as spheres. **e)** PCNA–DNA interface. PCNA interacting monomer is shown as a white ribbon, DNA primer and template strands are shown as white and purple sticks, respectively. Density around PCNA residues at a distance <4 Å from DNA phosphates is shown. Interacting residues are shown as sticks.

The B-form dsDNA exiting Pol κ core bends by ∼30°, and threads the PCNA ring with a ∼12° tilting angle (Figure 1f). The degree of tilting of DNA traversing PCNA is slightly larger than that observed in the processive Pol δ holoenzyme (∼4°), but similar to that observed in the two Pol δ conformers where PCNA is tilted (∼10°)^41^. Conversely, the pattern of interactions is different from those observed in all Pol δ conformers^41^, and mainly involves the PCNA protomer bound to Pol κ (Figure 3d). Clear density is observed for the side chains of Lys20 and Lys80 establishing electrostatic contacts with the phosphate of nucleotide 9 in the primer strand, and those of Asn84 and Arg210 interacting with phosphates of nucleotides 22 and 13 in the template strand, respectively (Figure 3e). These contacts are expected to be weak considering the very low affinity of the PCNA–DNA interaction (*K*_*d*_ ∼ 0.7 mM)^43^, and the distinct combination of PCNA residues interacting with DNA in different structures^41,43,44^. The PCNA inner surface therefore provides a flexible electrostatic screen for the DNA to pass through unhindered, and can adapt to different directions of the duplex DNA leaving the polymerase active site (Figure 3d).

### MD simulations of Pol κ holoenzyme in the absence of DNA

The C-terminal region of Pol κ PAD domain is the only site of binding to PCNA (Figure 2a-c) and does not participate in any interaction with DNA. The small surface of the PIP-box interaction raises the possibility that, in the absence of DNA, Pol κ bound to PCNA may sample conformations different from that in the holoenzyme. Indeed, flexible binding to the sliding clamp in the apo form was previously suggested for Y-family polymerases Pol IV^45^ and Dpo4^46^. We explored this scenario by performing Molecular Dynamics (MD) simulations of Pol κ–PCNA complex based on the Cryo-EM structure after removing the DNA (apo1 model) and on the same model but with Pol κ core in the orientation as in the X-ray structure of Pol κ apo form^33^ (apo2 model) (Figure 1b). Two 400-ns simulations for each starting model were performed; a time scale that does not allow equilibrated sampling of the conformational space but can probe flexibility and fast transitioning among potential conformations. Principal component analysis of the MD trajectories shows that the models sample wide yet different conformational space, indicative of large-scale conformational changes (Figure 4a; Supplementary Figure 6). Across all simulations, Pol κ maintains the PIP-box interaction with PCNA but displays high inter-molecular and inter-domain conformational flexibility due to two flexible hinges, one connecting the PIP-box to the PAD domain and centered on the inverting helix, and one connecting the PAD to the core domain (Figure 4b and Supplementary Movie 1-4). While Pol κ overall conformation fluctuates as its domains move around the two hinge regions, Pol κ individual domains and PCNA do not show significant variations, apart from minor shifts in Pol κ fingers subdomain (Supplementary Movie 1-4). Taken together, the Cryo-EM structure and MD simulations suggest that Pol κ bound to PCNA is able to switch from a flexible “carrier state”, characterized by high conformational freedom of the core domain relative to the PAD domain and the PAD domain relative to PCNA, to a rigid “active state” engaged for DNA synthesis (Figure 1g).

**Figure 4.**
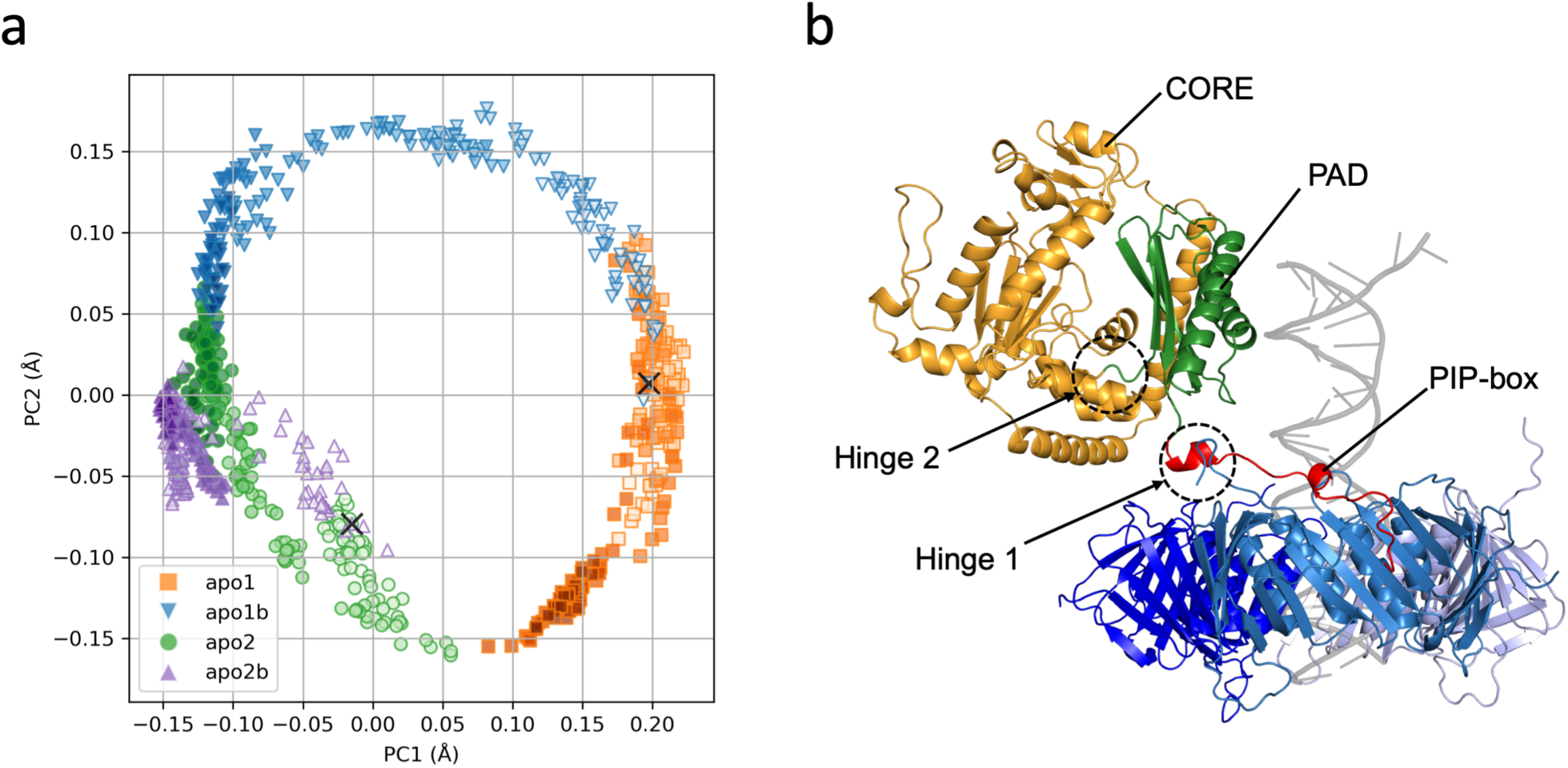
MD simulations of Pol κ–PCNA complex. **a)** Plot of the projection of the MD trajectories of the Pol κ–PCNA complex onto the first two principal components starting from model apo1 (orange squares and blue triangles) or apo2 (green circles and purple triangles). Intensity of symbol colors is proportional to the time evolution of the trajectories. The cross symbols represent the position at the start of the simulations. **b)** Structure of one frame of the apo1 MD trajectory, showing the large changes in the relative orientation of the core and PAD domains of Pol κ and PCNA. Pol κ core and PAD domains are shown as orange and green ribbons, respectively. PCNA protomers are shown as ribbons in different shades of blue. The DNA in the position corresponding to Pol κ holoenzyme is shown as transparent grey ribbon. The locations of the two hinge regions conferring flexibility to the complex are indicated.

### Cryo-EM structure of stalling Pol δ holoenzyme

We captured a non-replicating Pol δ holoenzyme by vitrifying a solution containing Pol δ heterotetramer, PCNA, a (25/38) P/T DNA substrate and a mixture of deoxynucleotides that allows up to 6-nt elongation of the primer, beyond which synthesis should stall for the lack of the required pairing nucleotide (Figure 5a). We obtained a reconstruction of the complex at 4.7 Å resolution and built a model of the four Pol δ subunits, P/T DNA and PCNA guided by the 3.0 Å-resolution structure of the processive Pol δ holoenzyme^41^ (Figure 5b-i, Supplementary Figures 7-8; Supplementary Table 1). The overall architecture is analogous to that of processive Pol δ^41^, displaying Pol δ anchored to one PCNA protomer through the PIP-box in the C-terminal domain of the p125 subunit (CTD), the regulatory subunits positioned sideways, and DNA threaded through PCNA almost perpendicularly to the ring plane (Figure 5f). However, the position of the DNA relative to the Pol δ catalytic subunit is different and conformational changes in Pol δ fingers and thumb subdomains as well as in the regulatory subunits are observed (Figure 6a-b; Supplementary Movie 5).

**Figure 5.**
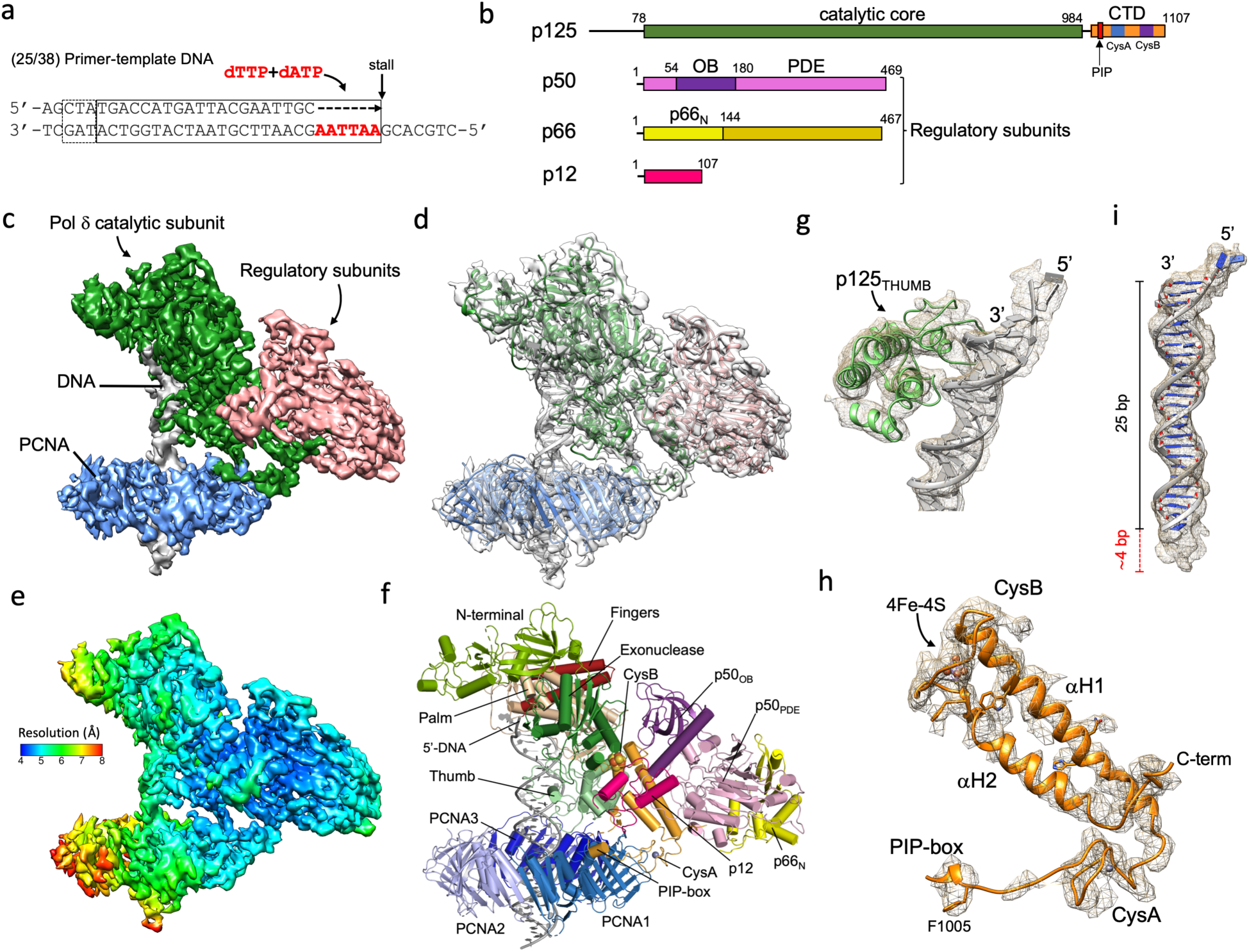
Cryo-EM structure of the stalled Pol δ–DNA–PCNA complex. **a)** Sequence of the DNA P/T substrate and dNTPs to generate a stalled complex. The solid box includes the base pairs that were modelled, the dotted box the base pairs accounted for the extended Cryo-EM density at the upstream region of dsDNA exiting PCNA, shown in panel i). **b)** Domain organization of human Pol δ; CTD: C-terminal domain; OB: oligonucleotide binding domain; PDE: phosphodiesterase domain; PIP: PCNA-interacting motif. **c)** Cryo-EM density map of stalled Pol δ holoenzyme complex colored by components. **d)** Stalled Pol δ holoenzyme structure fitted into the Cryo-EM map. **e)** Cryo-EM map of stalled Pol δ holoenzyme color-coded by local resolution. **f)** Structure of stalled Pol δ holoenzyme colored by domain. **g)** Map region around the thumb domain of Pol δ p125 subunit (green ribbon) and P/T DNA (grey ribbon). **h)** Map region around the CTD of Pol δ p125 subunit (orange ribbon). The iron sulfur cluster (4F–4S) and some of the bulky side chains are shown as sticks, the zinc atom in CysA as a sphere. **i)** Map region around the DNA substrate at a lower contour level, showing the extended density at the upstream dsDNA region.

**Figure 6.**
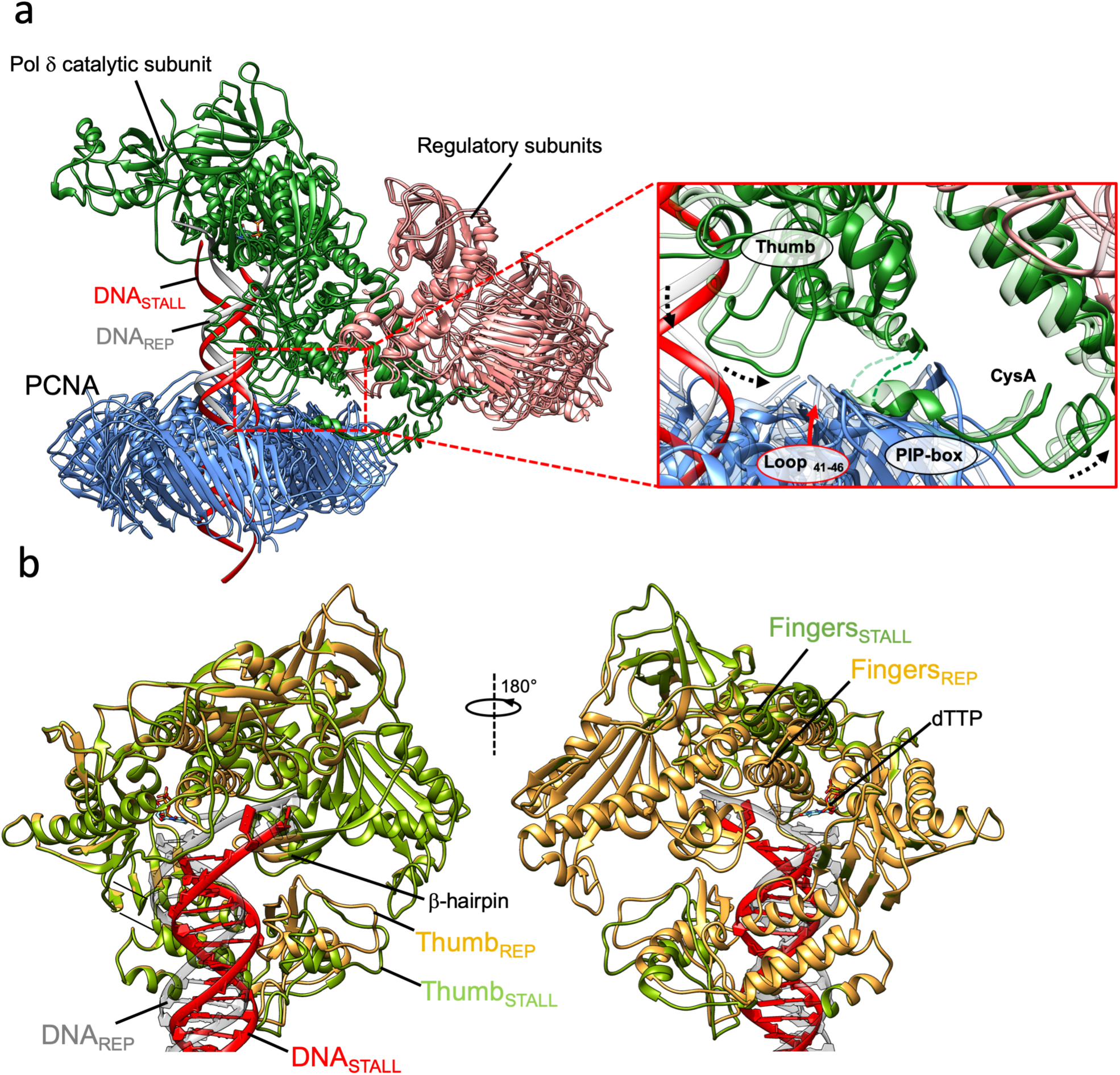
(a) Comparison of stalled and replicating Pol δ holoenzymes. **a)** Stalled and replicating Pol δ–PCNA–DNA complexes aligned on the catalytic subunit. DNA in the stalled (STALL) and replicating (REP) complex is shown as a red and gray ribbon, respectively. The poor alignment of the regulatory subunits and PCNA denotes the conformational changes induced by the release of DNA from the active site. The inset shows a close-up of the Pol δ–PCNA interface with the stalled and replicating structures as solid and transparent ribbons, respectively. The dotted arrows indicate the domain shifts induced by DNA release. The thumb outward displacement pulls the PCNA loop spanning residues 41-46 and this shift is propagated to the CTD and regulatory subunits of Pol δ. **b)** Overlay of the catalytic domain of Pol δ in the stalled (green) and replicating (orange) state in ribbon representation, highlighting the different conformation of DNA, fingers and thumb subdomains. The incoming nucleotide (dTTP) in the replicating state is shown as sticks. The DNA in the stalled and replicating state is colored red and grey, respectively.

No density of an incoming nucleotide is detected and the fingers subdomain is in the “open” conformation (Figure 6b and Supplementary Figure 9). Compared to the processive complex, the P/T DNA undergoes a concerted rotation and downward shift which moves it out of the active site, with a coordinated outward displacement of the polymerase thumb subdomain (Figure 6b). The thumb pulls a loop on PCNA front face inducing a tilt in the clamp ring and this movement propagates to the CTD and regulatory subunits (Figure 6a). Interestingly, the interaction between the thumb and the PCNA loop spanning residues 41-46 has been previously shown to regulate the replication activity of Pol δ holoenzyme^41^. While density connecting the thumb and the PCNA loop is observed in the map of the stalled holoenzyme, the map resolution prevented the assignment of the residues mediating the interaction. The change in the relative orientation of the p125 and regulatory subunits under stalling conditions highlights the overall plasticity of Pol δ holoenzyme, which has been previously described in yeast Pol δ in the absence of PCNA^47^.

The map density at the 5’ end of the template strand accounts for 2 nucleotides which project away from the catalytic core (Figure 5g and i). Compared to the replicative structure, the shift of the unpaired segment at the 5’ end of the template (Figure 6a-b) disrupts the interactions with the β-hairpin in the exonuclease domain spanning residues 431-449, and density at the tip of the β-hairpin is missing (residues 437-442), suggesting that the β-hairpin becomes flexible (Figure 6b). We could model 26 bp of the P/T DNA substrate (Figure 5a), yet density at the upstream end of dsDNA exiting the PCNA hole extends for further ∼3 bp when the map is inspected at a lower contour level (Figure 5i), demonstrating that Pol δ has extended the primer. Thus, in the absence of an incoming nucleotide, the processed DNA substrate is released from the active site through a conformational change of the thumb domain, but is retained to the complex. Opening of the thumb domain in a non-polymerizing binary complex of yeast Pol α and P/T DNA was previously observed, and postulated to be at the basis of a release mechanism of the P/T DNA^48^. Interestingly, the DNA is more exposed in the non-replicating versus processive complex, and is held in place primarily through its interaction with the thumb domain (Figure 6b). As discussed below, the release of P/T DNA from the active site may facilitate its handoff to a TLS polymerase.

## Discussion

### Functional implications of Pol κ holoenzyme structure

The structure of Pol κ holoenzyme is, to our knowledge, the first reported near-atomic resolution structure of a Y-family DNA polymerase bound to its processivity factor and DNA. The most striking feature consists in the sharply angled orientation of Pol κ core relative to the PCNA ring, and the resulting bending of dsDNA threading the PCNA central hole. The interaction tethering Pol κ to PCNA in this tilted position is mediated by the internal PIP-box (^526^QRSIIGFL) adjacent to the PAD domain of Pol κ, while the PIP-box (^862^KHTLDIFF) at the extreme C-terminus does not participate in the interaction. In agreement, previous studies showed that mutation of Pol κ residues Phe532 and Ile533 to alanines impairs stimulation of DNA synthesis of Pol κ by PCNA, while mutation of Phe868 and Phe869 has no effect on Pol κ activity^20^. The long region C-terminal to the internal PIP-box (residues 535-870) is invisible in the Cryo-EM map, suggesting it is largely disordered. In Pol η, flexibility of the C-terminal region has been previously observed experimentally^49^, and appears as a common characteristic of eukaryotic Y-family polymerases^50^. The disordered C-terminal region of Pol κ contains two UBZ zinc fingers (Figure 1a). Both UBZs bear a striking sequence similarity with Rad18-UBZ^51^, which binds ubiquitin with micromolar affinity^51^. In agreement with a current model of function, Pol κ UBZs may bind the ubiquitin moieties located at the back face of PCNA mono-ubiquitylated at K164 by Rad6–Rad18^52^, aiding the recruitment of Pol κ to sites of damage^37–39^. Pol κ holoenzyme structure is compatible with this model, since the long flexible C-terminus of Pol κ may easily bring the UBZs in proximity to one or more ubiquitins attached to the PCNA homotrimer (Figure 7a). In the human system, this model has been recently challenged by *in vitro* experiments using primer extension assays by Pol δ showing that the activity of Pol η and κ is independent of PCNA ubiquitylation^31,53,54^. *In vivo*, however, the secondary interaction mediated by ubiquitin may help Pol η or Pol κ to outcompete other proteins present at the replication fork, such as FEN1, Lig1 and PAF15, which all bind PCNA via similar PIP-box interactions^55–57^. In fact, conflicting results on the role of PCNA ubiquitylation is also observed in yeast using primer extension assays by Pol δ^58,59^. Nonetheless, a recent study showed that PCNA ubiquitination stimulates the recruitement of Pol η in a fully reconstituted yeast replisome, underlying the importance of studying the role of PCNA ubiquitination in TLS within the context of the replisome^12^. Because PCNA ubiquitylation exerts only a marginal effect in stimulation of DNA synthesis by Pol κ^20^, it is likely that the recruitment of Pol κ to PCNA principally functions through binding of the PIP-box adjacent to the PAD domain, as observed in the Cryo-EM structure.

**Figure 7.**
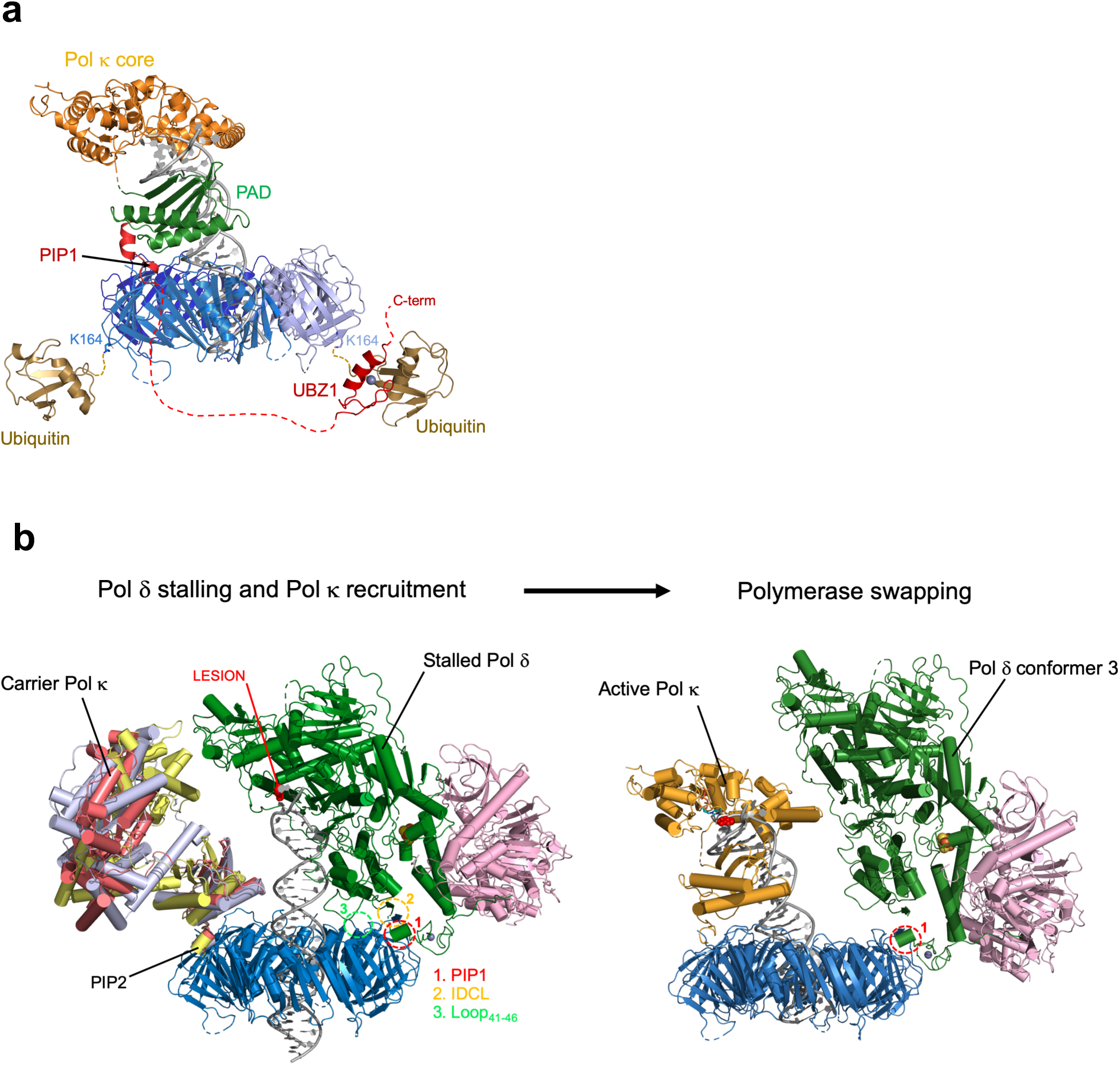
Proposed models of function of Pol κ holoenzyme in TLS. **a)** Pol κ holoenzyme interaction with ubiquitylated PCNA. Pol κ binding to PCNA brings Pol κ UBZ zinc fingers in proximity to the ubiquitin molecules at the back face of PCNA mono-ubiquitylated at K164 (PDB ID 3TBL)^92^. For simplicity, only UBZ1 was modelled. The long red dotted line represents the Pol κ disordered region spanning residues 534-617. The homology model for Pol κ UBZ1/ubiquitin complex was built with HHpred^93^, based on the NMR structure of Rad18-UBZ/ubiquitin complex (PDB ID 2MRE)^51^ **b)** PCNA-directed polymerase swapping in TLS. At a lesion on the DNA template strand (highlighted in red), Pol δ holoenzyme stalls (structure of stalled Pol δ in this study), DNA is released from the active site but remains associated to the complex. Pol κ is recruited to the exposed PIP-box site in the “carrier state” characterized by conformational flexibility of the PAD and core domains. Pol κ conformational sampling displaces Pol δ which tilts (Conformer 3; PDB ID 6S1O)^41^, further unmasking P/T DNA for its final transfer to Pol κ active site (Pol κ holoenzyme structure in this study). Tilting of Pol δ is achieved by disruption of 2 of the 3 indicated contact points with PCNA^41^, and allows to accommodate actively synthesizing Pol κ on PCNA in the “active state” without steric clashes. It is possible that Pol δ dissociates into solution prior to or during the formation of the Pol κ’s active state. In the left panel, Pol κ overlaid structures correspond to three frames from the MD simulation starting from apo1 model.

Therefore, the PAD domain of Pol κ docks the polymerase to PCNA, driving the assembly of the holoenzyme. In absence of DNA, our MD simulations predict that Pol κ bound to PCNA samples a wide conformational space, due to the conformational freedom of the core and PAD domains which results in an ensemble of different orientations of Pol κ relative to PCNA (carrier state). Binding to P/T DNA locks the polymerase–clamp complex into a rigid conformation that is competent for catalysis (active state). Thus, PCNA must function to recruit the “carrier state” polymerase to damaged P/T junctions in at least two critical situations *in vivo*: when the polymerase is located away from the target P/T junction, and when the junction is transiently engaged by another stalled polymerase. In the first instance, binding of Pol κ PAD domain to PCNA encircling dsDNA, and the disengagment of Pol κ core from the DNA, would ensure a rapid relocation of the polymerase to the target P/T junction due to the fast 1D diffusion of PCNA on dsDNA (diffusion coefficient ∼ 1 μm^2^s^−1^)^60,61^. The second situation pertains to one of the key steps of the TLS mechanism, namely the switch from the stalled replicative to the TLS polymerase to restart synthesis. As discussed below, the structures of Pol κ and stalled Pol δ holoenzymes may contribute to a better understanding of this molecular process.

### Model for PCNA-directed polymerase swapping in TLS

Eukaryotic Pol δ bound to PCNA replicates the DNA lagging strand, and is also critical for recoupling of leading-strand synthesis to the CMG helicase following lesion bypass^12^, but its synthetic activity and fidelity are impaired by damaged templates, particularly containing bulky lesions or abasic sites^62^. How a damaged DNA template may be transferred from a stalled Pol δ to a TLS polymerase to restart synthesis is unclear. Biochemical experiments showed that the higher affinity of Pol η for P/T DNA relative to Pol δ drives the exchange of the two polymerases in human TLS, and that PCNA is retained on the DNA substrate during the competition^53^. This agrees with a recent live-cell imaging analysis showing that, for the length of time PCNA is retained on damaged DNA, Pol δ and a TLS polymerase could have exchanged around 60 times^63^.

In the Cryo-EM structure of the non-replicating Pol δ holoenzyme reported herein, the DNA is released from the active site due to the absence of the required pairing nucleotide, but remains attached to the thumb domain of the polymerase. This structure may therefore represent a stalling state prior to the handoff of DNA to a TLS polymerase, and implies that Pol δ has to depart from DNA for the handoff to occur. Based on the stalled Pol δ structure, the structure of Pol κ holoenzyme and MD simulations, and the previously determined structures of tilted conformations of Pol δ bound to PCNA^41^, we propose a model for the steps of polymerase swapping in TLS (Figure 7b). At a DNA lesion, Pol δ stalls and releases P/T DNA from the active site. Pol κ docks to the vacating most exposed PCNA protomer in the flexible “carrier state”. At this point, either conformational sampling of Pol κ may actively displace Pol δ from P/T DNA, or Pol δ may autonomously depart from the DNA, allowing DNA handoff and the final assembly of Pol κ holoenzyme. In both cases, full removal of Pol δ from PCNA is not necessary, as Pol δ may loose the interactions with the IDCL and Loop_41-46_ of PCNA and retain the PIP-box interaction, resulting in a PCNA tilting which provides enough room to accomodate actively synthesising Pol κ without steric clash. In fact, Pol δ in the tilted conformers is bound to PCNA only via the PIP-box^41^, suggesting that in the absence of DNA it will be flexibly tethered to PCNA and sampling different conformations. Once the lesion is bypassed, the DNA may be trasferred back to Pol δ by reversing the order of the described steps.

While this mechanism is compatible with the “toolbelt model” in TLS^64^, it also accounts for the possibility that Pol δ may fully dissociate from PCNA and reassociate after the TLS polymerase has bypassed the lesion, as previously suggested in the case of human Pol η^53,54^. On the other hand, we find it unlikely that the TLS polymerase may compete with Pol δ for binding to the same PIP-box site on PCNA, considering the very high affinity of Pol δ for PCNA encircling DNA (*K*_*d*_ < 10 nM)^54,65^ and the comparetively low affinities of human Y-family TLS polymerases for PCNA (*K*_*d*_ > 100 nM)^53,66^. Interestingly, biochemical experiments showed the coexistence of Pol III and Pol IV on β-clamp in bacterial TLS^67,68^, and a supra-holoenzyme consisting of PolB1 and PolY simultaneously bound to PCNA has been characterized in archeal TLS^69^. If a “TLS toolbelt” involving Pol δ exists in eukaryotes remains to be determined. Our previously reported structure of human Pol δ and FEN1 simultaneously bound to PCNA^41^ is the first direct evidence of a toolbelt in eukaryotes. It is therefore possible that a TLS polymerase may replace FEN1 on PCNA to perform lesion bypass. Ubiquitination of PCNA may help recruit Pol κ allowing it to outcompete FEN1. Our structures show that the coexistence of Pol δ and Pol κ on a PCNA ring would be sterically allowed through the tilted conformation of PCNA, which is supported by the intrinsic flexibility of the PIP-box interaction with the C-terminal domain of the p125 subunit of Pol δ.

## Materials and Methods

### Protein Expression and Purification

Human Pol κ was purified using a modified version of previously published protocol^70^. *E. coli* codon-optimized sequence of human full length Pol κ (accession no. NP057302) was cloned into a pE-SUMOpro expression vector (Lifesensors) using Gibson assembly technology. The Pol κ plasmid was transformed into *E. coli* strain BL21 (DE3) competent cells (Novagen) that were grown on agar plates containing 50 μg/ml kanamycin. Several colonies were randomly selected and checked for expression level. Pol κ was overexpressed by growing the transformed cells into 10 L of 2YT media (Teknova) supplemented with kanamycin. Cells were incubated at 24 °C till the OD_600_ reached 0.8 and then protein expression was induced at 0.1 mM isopropyl β-D-thiogalactopyranoside (IPTG) concentration. The cells were incubated further for 19 hrs at 16 °C, harvested by centrifugation at 5,500 xg for 10 min then re-suspended in lysis buffer [50 mM Tris pH (8), 750 mM NaCl, 40 mM Imidazole, 5 mM β-Mercaptoethanol, 0.2% NP-40, 1 mM PMSF, 5% Glycerol and EDTA free protease inhibitor cocktail tablet/50ml (Roche, UK)]. All subsequent steps were performed at 4 °C. The cells were lysed by 1 hour incubation on ice using lysozyme at final concentration of 2 mg/ml followed by mechanical disruption by sonication. Cell debris was then removed by centrifugation at 22,040 xg for 30 min. The decanted supernatant was directly loaded onto HisTrap HP 5 ml affinity column (GE Healthcare) pre-equilibrated with Buffer A [50 mM Tris (pH 7.5), 500 mM NaCl, 40 mM Imidazole, 5 mM β-Mercaptoethanol and 5% Glycerol]. The column was then washed with 10 column volumes (CVs) of Buffer A and eluted by 10 CV gradient against Buffer B [50 mM Tris (pH 7.5), 500 mM NaCl, 500 mM Imidazole, 5 mM β-Mercaptoethanol and 5% Glycerol]. The protein was eluted around 210 mM imidazole concentration. The fractions containing Pol κ were checked by SDS-PAGE. The peak fractions were then pooled and dialyzed overnight against dialysis buffer [50mM Tris (pH 7.5), 500 mM NaCl, 5 mM β-Mercaptoethanol and 5% Glycerol] containing SUMO protease (LifeSensors) in order to cleave the SUMO tag and generate Pol κ in the native form. The dialyzed sample was then loaded again onto HisTrap HP 5ml column using Buffer A and the native protein was collected in the flow-through fractions. Fractions that contained Pol κ were pooled, concentrated and then loaded onto HiLoad 16/600 Superdex 200 pg (GE Healthcare) equilibrated with storage buffer [50 mM Tris (pH 7.5), 300 mM NaCl and 1 mM DTT]. Fractions containing Pol κ were checked for purity, concentrated, flash frozen and stored at −80 °C.

Human Pol δ was cloned and expressed as described previously^41^. Briefly, all four Pol δ subunits encoding p125 (accession no. NP02682), p50 (accession no. NP006221), p66 (accession no. NP006582) and p12 (accession no. NP066996) were amplified and cloned to construct a single MultiBac™ expression plasmid. Bacmid DNA was generated by transforming single recombinant transfer vector encoding all four Pol δ subunits into DH10MultiBac™ cells. To prepare the baculovirus, bacmid DNA containing all four subunits was transfected into Sf9 cells using FuGENE® HD (Promega) per manufacturer’s instructions. The resulting baculovirus was then amplified twice to obtain a higher titer virus preparation (P3 virus). The expression of Pol δ then proceeded by transfecting 4 L of Sf9 suspension culture grown at a density of 2 × 10^6^ cells/mL with P3 virus for 66-72 hrs. Pol δ cleared lysate was then loaded onto a HisTrap column and eluted with low salt, followed by ion exchange chromatography on a Mono Q column, and finally size exclusion chromatography on HiLoad 16/600 Superdex 200 pg. Pure protein fractions were pooled, flash frozen and stored at −80°C.

Human PCNA used for the Pol κ replication assays was produced as described previously^41^. Briefly, full length human PCNA (accession no. NM182649) was cloned into pETDuet-1 MCS1 (Novagen) Amp^+^ to obtain 6× His N-terminally tagged protein, transformed into *E. coli* strain BL21 (DE3) cells and grown at 37 °C in 2YT media supplemented with ampicillin till reaching OD_600_ of 1.2. Protein expression was induced with 0.5 mM IPTG for 19 hrs at 16 °C. Cells were then harvested, and lysed with lysozyme followed by sonication. The cleared lysate was loaded onto a HisTrap column (GE Healthcare) and eluted with low salt, followed by anion exchange on a HiTrap Q column (GE Healthcare), and finally size exclusion chromatography on a HiLoad 16/600 Superdex 200 pg. Pure protein fractions were pooled, flash frozen, and stored at −80 °C. Recombinant PCNA used for the Cryo-EM study was produced as described previously^43^.

### DNA substrates

DNA oligos for the primer extension assays were synthesized and HPLC purified by IDT and The Midland Co. The non-damage substrate consisted of a 63 nt template with a biotin moiety attached to triethylene glycol (TEG) spacer at each end and a 28 nt primer whose 5’ end was labeled with Cy5. The sequences of both oligos, respectively, are as follows:/5′BiotinTEG/ATCTTCCTTCAACCAGCTTACCTTCAACGATTTAGGTTACCTTCAATGTCATGCTCGCGCTGA/3’BioTEG/) and /5’Cy5/CAGCGCGAGCATGACATTGAAGGTAACC-3’). The abasic substrate consisted of 55 nt template strand containing abasic site and 19 nt primer labeled with Cy5 at the 5’ end. The sequences of both oligos, respectively, are: 5’-CCCTCTAGAGGCGCGCCGGACATGTAAT(abasic)AACCATGGGAGACCGGGTACC CCC -3′ and /5′Cy5/GGGGGTACCCGGTCTCCCA-3′. Substrates were annealed by mixing both template and primer at 1:1 molar ratio in TE-100 buffer [50 mM Tris-HCl (pH 8.0), 1 mM EDTA and 100 mM NaCl] and heating at 95 °C for 5 min followed by slow cooling down to room temperature. Substrates were PAGE purified to >90% purity using 10% non-denaturing polyacrylamide gel electrophoresis (Invitrogen). The biotin-labeled substrate was incubated with a 2-fold molar excess of neutravidin before the primer extension assay. Finally, the substrates were aliquoted and stored at −20 °C.

For the substrates used in Cryo-EM, a template strand (5’-CTGCACGAATTAAGCAATTCGTAATCATGGTCATAGCT-3’) was annealed to either an unmodified primer (5’-AGCTATGACCATGATTACGAATTGC-3’) to form the P/T substrate, or to a primer containing a dideoxycytosine at the 3’ end (5’-AGCTATGACCATGATTACGAATTG(DOC)-3’) to form the P/T(ddC) substrate. In both cases, the oligos were mixed in an equimolar ratio in the presence of 20 mM Tris-HCl (pH 7.5) and 25 mM NaCl. The oligos were then annealed by heating at 92 °C for 2 min followed by slow cooling down to room temperature. All three oligos were purchased from Sigma Aldrich.

### Primer extension assay

The primer extension activity assays of Pol δ and Pol κ were performed on either the non-damage or abasic substrate in 10 µl total reaction volume. Briefly, 20 nM non-damage or abasic substrates were incubated with proteins at 30 °C for 5 mins or 2 mins, respectively, in the reaction buffer [40 mM Tris-HCl (pH 7.8), 50 mM NaCl, 0.2 mg/ml BSA, 1 mM DTT, 5 mM MgCl_2_, 1 mM ATP, 0.1 mM of each deoxynucleotide (dATP, dTTP, dGTP, and dCTP)]. The reactions were terminated with the addition of 10 µl of stop buffer [50 mM EDTA, 95% Formamide and 1.5% Ficoll] followed by heating at 95 °C for 3 min and cooling down on ice for 2 min. All products were resolved on 12% polyacrylamide gels containing 8 M urea and visualized using Typhoon Trio fluorescence imager (GE Healthcare).

### Cryo-EM grid preparation and data collection

For the Pol κ holoenzyme dataset, a 40 μl inject containing 3.75 μM P/T(ddC) DNA, 3.78 μM Pol κ, 1.5 μM PCNA trimer and 20 μM TTP was loaded onto a Superdex 200 increase 3.2/300 column (GE Life Sciences) equilibrated with a buffer comprising [25 mM HEPES (pH 7.5), 100 mM K-Ac, 10 mM CaCl_2_, 0.02% NP-40, 0.4 mM Biotin and 1 mM DTT]. 3 μl of a fraction corresponding to the first peak (supplementary figure xx, fraction and lane 2) was used. For the Pol δ holoenzyme dataset, 4.2 μM P/T DNA, 1.02 μM Pol δ, 0.84 μM PCNA and 20 μM each of dATP and dTTP were mixed in a buffer comprising [25 mM HEPES pH 7.5, 100 mM K-Ac, 10 mM MgCl_2_, 0.02% NP-40, 0.4 mM Biotin and 1 mM DTT]. 3 μl of this was used for the grid. For both complexes, UltrAuFoil® R1.2/1.3 Au 300 grids were glow discharged for 5 min at 40 mA on a Quorum Gloqube glow-discharge unit, then covered with a layer of graphene oxide (Sigma) prior to application of sample. Once the sample was applied to the grid, it was blotted for 3 seconds and plunge frozen into liquid ethane using a Vitrobot Mark IV (FEI Thermo Fisher), which was set at 4 °C and 100% humidity. Cryo-EM data for both complexes were collected on a Thermo Fisher Scientific Titan Krios G3 transmission electron microscope at the Midlands Regional Cryo-EM Facility at the Leicester Institute of Structural and Chemical Biology. For the Pol κ dataset, electron micrographs were recorded using a K3 direct electron detector (Gatan Inc.) at a dose rate of 11 e-/pix/sec and a calibrated pixel size of 1.09 Å. The data were collected with EPU 2.3. Data were acquired with a defocus range between −1.5 and −0.7 μm, in 0.2 μm intervals. For the Pol δ dataset, electron micrographs were recorded using a K2 Summit direct electron detector (Gatan Inc.) at a dose rate of 5.8 e-/pix/sec and a calibrated pixel size of 1.4 Å. This dataset was collected using a Volta phase plate with EPU 1.9. Focusing was performed at every hole using a nominal value of −0.6 μm.

### Cryo-EM image processing

Pre-processing of the Pol κ dataset was performed in Relion-3.1^71^ as follows: movie stacks imported in super resolution mode, then corrected for beam-induced motion and then integrated using Relion’s own implementation, using a binning factor of 2. All frames were retained and a patch alignment of 5 × 5 was used. Contrast transfer function (CTF) parameters for each micrograph were estimated by CTFFIND-4.1^72^. Integrated movies were inspected with Relion-3.1 for further image processing (2714 movies). Particle picking was performed in an automated mode using the Laplacian-of-Gaussian (LoG) filter implemented in Relion-3.1. All further image processing was performed in Relion-3.1. Particle extraction was carried out from micrographs using a box size of 300 pixels (pixel size: 1.086 Å/pixel). An initial dataset of 2.3 × 10^6^ particles was cleaned by 2D classification followed by 3D classification with alignment. 3D refinement and a few rounds of polishing and per-particle CTF refinement yielded a 3.93 Å structure of the Pol κ-PCNA-DNA complex.

Pre-processing of the Pol δ complex was performed in Relion-3.0 as follows: movie stacks were corrected for beam-induced motion and then integrated using MotionCor2^73^. All frames were retained and a patch alignment of 4 × 4 was used. CTF parameters for each micrograph were estimated by Gctf^74^. A total of 4837 micrographs were processed, with 985 of them acquired with a stage tilted by 40° to improve the angular distribution of particles. Integrated movies were inspected with Relion-3.0 for further image processing. Particle picking was performed in an automated mode using the Laplacian-of-Gaussian (LoG) filter implemented in Relion-3.0. Particle extraction was carried out from micrographs using a box size of 280 pixels (pixel size: 1.4 Å/pixel). An initial dataset of 3.2×10^6^ particles was cleaned by 2D classification followed by two rounds of 3D classification with alignment and one round without alignment. 3D refinement and several rounds of polishing yielded a 4.72 Å structure of the Pol δ–PCNA–DNA complex.

### Molecular modelling

#### Model building of Pol κ holoenzyme

The X-ray structure of the catalytic domain of human Pol κ bound to P/T DNA and dTTP (PDB ID 2OH2)^32^, and the structure of PCNA homotrimer (from PDB ID 6TNZ)^33^ were rigid-body fitted into the Cryo-EM map. N-Clasp residues 21-45 of Pol κ X-ray structure^32^ were invisible in the map and were deleted from the model. The upstream 19 base pairs of B-form duplex DNA were built with Chimera^75^ and Coot^76^ and real-space refined with Coot. The region of Pol κ at the PAD C-terminus (residues 517-534) was built and refined with Coot. The entire model of Pol κ holoenzyme was subjected to real-space refinement in Phenix^77^ with the application of secondary structure and stereochemical constraints.

#### Model building of stalled Pol κ holoenzyme

The model was built based on the structure of the replicating Pol κ holoenzyme (PDB: 6TNY)^41^, which was partitioned into the following sub-structures: p125 catalytic core, p125 CTD, p50–p66_N_–p12 subcomplex, and PCNA trimer. These structures were individually rigid-body fitted into the Cryo-EM map of the stalled Pol δ complex and edited with Coot. The thumb and fingers subdomains of the p125 subunit were edited with Coot and rigid-body fitted into the map. The 26-bp B-form DNA was built with 3D-DART^78^ and UCSF Chimera, and rigid-body fitted into the Cryo-EM map. The 5’ overhang in the template strand of the DNA substrate was built and real-space fitted into the Cryo-EM map with Coot. Because the map resolution did not allow to discriminate the identity of the nucleotides in the DNA substrate, and because it was not possible to unambiguously define the number of nucleotides inserted in the primer strand by Pol δ before stalling, all nucleotides of the primer and template strands were arbitrarily assigned as thymidines and adenosines, respectively. The final models were validated using Phenix.

### MD simulations

Simulations were started using two different models of the Pol κ–PCNA complex. The first model (apo1) was generated from the Cryo-EM structure of Pol κ holoenzyme after removing the DNA. The second model (apo2) was obtained from apo1 with the following steps: the X-ray structure of apo Pol κ (PDB ID 1T94)^33^ was aligned to Pol κ PAD domain in apo1 model. Pol κ core domain of apo1 model was then extracted and aligned to the core domain of 1T94, and the loop connecting the core and PAD domains was rebuilt. In both apo1 and apo2 models, some of the disordered residues that could not be resolved in the Cryo-EM structure were reconstructed with Modeller^79,80^. The regions of Pol κ spanning residues 1-36 and 225-281, absent in the Cryo-EM structure, were not reconstructed because obtaining conformationally converged ensembles for long disordered regions is slow and because there is no evidence that these regions participate in contacts with PCNA. Residues Met225 and Gln281 are in close proximity and were joint by modelling their positions and that of the two bridging residues Gly226 and Leu280. We used the recently developed DES-Amber force field which aims to correctly describe protein-protein interactions^81^. The setup of the simulation closely followed the procedure described in^82^. Briefly, the initial structures were solvated in a dodecahedron box 1 nm away from initial protein atoms. The water model used was TIP4P-D also parametrized in conjunction with the protein force field^81^. Na^+^ and Cl^−^ ions were added to simulate a 100 mM NaCl solution. This structure was minimized and run in an NVT ensemble for 2 ns and in an NPT ensemble for additional 2 ns. These simulations had positional restrains of 1000 kJ mol^−1^ nm^−1^ to all non-hydrogen protein atoms to allow relaxation of the solvent. From these structures, two unrestrained molecular dynamics simulations for each of the two systems were started. Virtual sites in the setup of these simulations were used, which allowed a time step of 4 fs. All simulations were run with Gromacs 2019.4^83–85^. The analysis of the trajectories was carried out with in-house scripts using MDTraj^86^. All four trajectories were concatened into a single trajectory. Then all frames of this trajectory were superimposed onto its first frame and Cartesian coordinates of the backbone atoms were used to calculate the principal components with the implementation in scikit-learn library^87^. Plots were produced with the Matplotlib library^88^. Tools in ENCORE to evaluate the convergence of the simulations^89^ were used. For visualization of the trajectories, VMD^90^ and Pymol^91^ were used.

## Acknowledgements

This research was supported by King Abdullah University of Science and Technology through core funding (to S.M.H.), and by the Wellcome Trust (to A.D.B.). R.C. acknowledges funding from the MINECO (CTQ2016-78636-P) and to AGAUR, (2017 SGR 324). The MD project has been carried out using CSUC resources. We acknowledge The Midlands Regional Cryo-EM Facility at the Leicester Institute of Structural and Chemical Biology (LISCB), major funding from MRC (MC_PC_17136). We thank Christos Savva (LISCB, University of Leicester) for his help in Cryo-EM data collection and advice on data processing.

## Author Contributions

M. Tehseen purified Pol δ, Pol κ, and PCNA, confirmed their activities, and helped in initiating the project; M. Tehseen, M. Takahashi, and M.A.S. optimized and perfomed the TLS assays; C.L. prepared the cryo-EM samples; C.L., T.J.R., C.S. and A.D.B. analysed the cryo-EM data; C.L. and A.D.B. built and refined the molecular models. R.C. performed and analysed the MD simulations. S.M.H. and A.D.B. conceived the research and wrote the article. All authors discussed the results and commented on the manuscript.

## Author Information

The maps of the Pol κ holoenzyme and stalled Pol δ–DNA–PCNA complex have been deposited in the EMBD with accession codes EMD-11291 and EMD-11290, and the atomic models in the Protein Data Bank under accession codes PDB 6ZMH and 6ZMF. The authors declare no competing financial interests. Correspondence and requests for materials should be addressed to S.M.H. or A.D.B.

## References

1. Klarer, A. C. & McGregor, W. Replication of Damaged Genomes. Crit. Rev. Eukaryot. Gene Expr. 21, 323–336 (2011).

2. Huen, M. S. Y. & Chen, J. Assembly of checkpoint and repair machineries at DNA damage sites. Trends Biochem. Sci. 35, 101–108 (2010).

3. Yang, W. & Woodgate, R. What a difference a decade makes: Insights into translesion DNA synthesis. Proc. Natl. Acad. Sci. 104, 15591–15598 (2007).

4. McCulloch, S. D. & Kunkel, T. A. The fidelity of DNA synthesis by eukaryotic replicative and translesion synthesis polymerases. Cell Res. 18, 148–161 (2008).

5. Marians, K. J. Lesion Bypass and the Reactivation of Stalled Replication Forks. Annu. Rev. Biochem. 87, 217–238 (2018).

6. Budzowska, M. & Kanaar, R. Mechanisms of Dealing with DNA Damage-Induced Replication Problems. Cell Biochem. Biophys. 53, 17–31 (2009).

7. Lange, S. S., Takata, K. & Wood, R. D. DNA polymerases and cancer. Nat. Rev. Cancer 11, 96–110 (2011).

8. Yang, Y. et al. Diverse roles of RAD18 and Y-family DNA polymerases in tumorigenesis. Cell Cycle 17, 833–843 (2018).

9. Tonzi, P. & Huang, T. T. Role of Y-family translesion DNA polymerases in replication stress: Implications for new cancer therapeutic targets. DNA Repair (Amst). 78, 20–26 (2019).

10. Yang, W. & Gao, Y. Translesion and Repair DNA Polymerases: Diverse Structure and Mechanism. Annu. Rev. Biochem. 87, 239–261 (2018).

11. Maxwell, B. A. & Suo, Z. Recent insight into the kinetic mechanisms and conformational dynamics of Y-family DNA polymerases. Biochemistry 53, 2804–2814 (2014).

12. Guilliam, T. A. & Yeeles, J. T. P. Reconstitution of translesion synthesis reveals a mechanism of eukaryotic DNA replication restart. Nat. Struct. Mol. Biol. 27, 450–460 (2020).

13. Choe, K. N. & Moldovan, G.-L. Forging Ahead through Darkness: PCNA, Still the Principal Conductor at the Replication Fork. Mol. Cell 65, 380–92 (2017).

14. Vaisman, A. & Woodgate, R. Translesion DNA polymerases in eukaryotes: what makes them tick? Crit. Rev. Biochem. Mol. Biol. 52, 274–303 (2017).

15. Mondol, T., Stodola, J. L., Galletto, R. & Burgers, P. M. PCNA accelerates the nucleotide incorporation rate by DNA polymerase delta. Nucleic Acids Res. 47, 1977–1986 (2019).

16. Hedglin, M. & Benkovic, S. J. Regulation of Rad6/Rad18 Activity During DNA Damage Tolerance. Annu. Rev. Biophys. 44, 207–228 (2015).

17. Hoege, C., Pfander, B., Moldovan, G. L., Pyrowolakis, G. & Jentsch, S. RAD6-dependent DNA repair is linked to modification of PCNA by ubiquitin and SUMO. Nature 419, 135–141 (2002).

18. Haracska, L., Torres-Ramos, C. A., Johnson, R. E., Prakash, S. & Prakash, L. Opposing Effects of Ubiquitin Conjugation and SUMO Modification of PCNA on Replicational Bypass of DNA Lesions in Saccharomyces cerevisiae. Mol. Cell. Biol. 24, 4267–4274 (2004).

19. Sale, J. E., Lehmann, A. R. & Woodgate, R. Y-family DNA polymerases and their role in tolerance of cellular DNA damage. Nat. Rev. Mol. Cell Biol. 13, 141–152 (2012).

20. Masuda, Y. et al. Different types of interaction between PCNA and PIP boxes contribute to distinct cellular functions of Y-family DNA polymerases. Nucleic Acids Res. 43, 7898–7910 (2015).

21. Plosky, B. S. et al. Controlling the subcellular localization of DNA polymerases ι and η via interactions with ubiquitin. EMBO J. 25, 2847–2855 (2006).

22. Bienko, M. et al. Ubiquitin-Binding Domains in Y-Family Polymerases Regulate Translesion Synthesis. Science (80-.). 310, 1821–1824 (2005).

23. Despras, E., Delrieu, N., Garandeau, C., Ahmed-Seghir, S. & Kannouche, P. Regulation of the Specialized DNA Polymerase Eta: Revisiting the Biological Relevance of Its PCNA- and Ubiquitin-Binding Motifs. Environ. Mol. Mutagen. 53, 752–65 (2012).

24. Kannouche, P. L., Wing, J. & Lehmann, A. R. Interaction of human DNA polymerase η with monoubiquitinated PCNA: A possible mechanism for the polymerase switch in response to DNA damage. Mol. Cell 14, 491–500 (2004).

25. Watanabe, K. et al. Rad18 guides polη to replication stalling sites through physical interaction and PCNA monoubiquitination. EMBO J. 23, 3886–3896 (2004).

26. Masuda, Y., Piao, J. & Kamiya, K. DNA Replication-Coupled PCNA Mono-Ubiquitination and Polymerase Switching in a Human In Vitro System. J. Mol. Biol. 396, 487–500 (2010).

27. Prakash, S., Johnson, R. E. & Prakash, L. Eukaryotic Translesion Synthesis DNA Polymerases: Specificity of Structure and Function. Annu. Rev. Biochem. 74, 317–353 (2005).

28. Stern, H. R., Sefcikova, J., Chaparro, V. E. & Beuning, P. J. Mammalian DNA Polymerase Kappa Activity and Specificity. Molecules 24, 2805 (2019).

29. Washington, M. T., Johnson, R. E., Prakash, L. & Prakash, S. Human DINB1-encoded DNA polymerase κ is a promiscuous extender of mispaired primer termini. Proc. Natl. Acad. Sci. U. S. A. 99, 1910–1914 (2002).

30. Wolfle, W. T., Washington, M. T., Prakash, L. & Prakash, S. Human DNA polymerase κ uses template-primer misalignment as a novel means for extending mispaired termini and for generating single-base deletions. Genes Dev. 17, 2191–2199 (2003).

31. Barnes, R. P., Hile, S. E., Lee, M. Y. & Eckert, K. A. DNA polymerases eta and kappa exchange with the polymerase delta holoenzyme to complete common fragile site synthesis. DNA Repair (Amst). 57, 1–11 (2017).

32. Lone, S. et al. Human DNA Polymerase κ Encircles DNA: Implications for Mismatch Extension and Lesion Bypass. Mol. Cell 25, 601–614 (2007).

33. Uljon, S. N. et al. Crystal structure of the catalytic core of human DNA polymerase kappa. Structure 12, 1395–1404 (2004).

34. Wong, J. H., Fiala, K. A., Suo, Z. & Ling, H. Snapshots of a Y-Family DNA Polymerase in Replication: Substrate-induced Conformational Transitions and Implications for Fidelity of Dpo4. J. Mol. Biol. 379, 317–330 (2008).

35. Trincao, J. et al. Structure of the Catalytic Core of S. cerevisiae DNA polymerase η: Implications for translesion DNA synthesis. Mol. Cell 8, 417–426 (2001).

36. Silverstein, T. D. et al. Structural basis for the suppression of skin cancers by DNA polymerase eta. Nature 465, 1039–1044 (2010).

37. Bi, X. et al. Rad18 Regulates DNA Polymerase κ and Is Required for Recovery from S-Phase Checkpoint-Mediated Arrest. Mol. Cell. Biol. 26, 3527–3540 (2006).

38. Guo, C., Tang, T. S., Bienko, M., Dikic, I. & Friedberg, E. C. Requirements for the interaction of mouse Polκ with ubiquitin and its biological significance. J. Biol. Chem. 283, 4658–4664 (2008).

39. Jones, M. J. K., Colnaghi, L. & Huang, T. T. Dysregulation of DNA polymerase κ recruitment to replication forks results in genomic instability. EMBO J. 31, 908–918 (2012).

40. Lee, M. Y. W. T., Wang, X., Zhang, S., Zhang, Z. & Lee, E. Y. C. Regulation and modulation of human DNA polymerase δ activity and function. Genes (Basel). 8, 190 (2017).

41. Lancey, C. et al. Structure of the processive human Pol d holoenzyme. Nat. Commun. 11, 1109 (2020).

42. Boehm, E. M. & Washington, M. T. R.I.P. to the PIP: PCNA-binding motif no longer considered specific. BioEssays 38, 1117–1122 (2016).

43. De March, M. et al. Structural basis of human PCNA sliding on DNA. Nat. Commun. 8, 13935 (2017).

44. McNally, R., Bowman, G. D., Goedken, E. R., O’Donnell, M. & Kuriyan, J. Analysis of the role of PCNA-DNA contacts during clamp loading. BMC Struct. Biol. 10, 3 (2010).

45. Bunting, K. A., Roe, S. M. & Pearl, L. H. Structural basis for recruitment of translesion DNA polymerase Pol IV/DinB to the beta-clamp. EMBO J. 22, 5883–5892 (2003).

46. Xing, G., Kirouac, K., Shin, Y. J., Bell, S. D. & Ling, H. Structural insight into recruitment of translesion DNA polymerase Dpo4 to sliding clamp PCNA. Mol. Microbiol. 71, 678–691 (2009).

47. Jain, R. et al. Cryo-EM structure and dynamics of eukaryotic DNA polymerase δ holoenzyme. Nat. Struct. Mol. Biol. 26, 955–962 (2019).

48. Perera, R. L. et al. Mechanism for priming DNA synthesis by yeast DNA Polymerase α. Elife 2, e00482 (2013).

49. Powers, K. T., Elcock, A. H. & Washington, M. T. The C-terminal region of translesion synthesis DNA polymerase η is partially unstructured and has high conformational flexibility. Nucleic Acids Res. 46, 2107–2120 (2018).

50. Ohmori, H., Hanafusa, T., Ohashi, E. & Vaziri, C. Separate roles of structured and unstructured regions of Y-family DNA polymerases. Advances in protein chemistry and structural biology 78, (Elsevier, 2009).

51. Rizzo, A. A., Salerno, P. E., Bezsonova, I. & Korzhnev, D. M. NMR Structure of the Human Rad18 Zinc Finger in Complex with Ubiquitin Defines a Class of UBZ Domains in Proteins Linked to the DNA Damage Response. Biochemistry 53, 5895–5906 (2014).

52. Freudenthal, B. D., Gakhar, L., Ramaswamy, S. & Washington, M. T. Structure of monoubiquitinated PCNA and implications for translesion synthesis and DNA polymerase exchange. Nat. Struct. Mol. Biol. 17, 479–484 (2010).

53. Hedglin, M., Pandey, B. & Benkovic, S. J. Characterization of human translesion DNA synthesis across a UV-induced DNA lesion. Elife 5, e19788 (2016).

54. Hedglin, M., Pandey, B. & Benkovic, S. J. Stability of the human polymerase δ holoenzyme and its implications in lagging strand DNA synthesis. Proc. Natl. Acad. Sci. 113, E1777–86 (2016).

55. Sakurai, S. et al. Structural basis for recruitment of human flap endonuclease 1 to PCNA. EMBO J. 24, 683–693 (2005).

56. Montecucco, A. et al. DNA ligase I is recruited to sites of DNA replication by an interaction with proliferating cell nuclear antigen: Identification of a common targeting mechanism for the assembly of replication factories. EMBO J. 17, 3786–3795 (1998).

57. De Biasio, A. et al. Structure of p15PAF-PCNA complex and implications for clamp sliding during DNA replication and repair. Nat. Commun. 6, 6439 (2015).

58. Garg, P. & Burgers, P. M. Ubiquitinated proliferating cell nuclear antigen activates translesion DNA polymerases eta and REV1. Proc. Natl. Acad. Sci. 102, 18361–18366 (2005).

59. Haracska, L., Unk, I., Prakash, L. & Prakash, S. Ubiquitylation of yeast proliferating cell nuclear antigen and its implications for translesion DNA synthesis. Proc. Natl. Acad. Sci. U. S. A. 103, 6477–6482 (2006).

60. Kochaniak, A. B. et al. Proliferating Cell Nuclear Antigen Uses Two Distinct Modes to Move along DNA * □. 284, 17700–17710 (2009).

61. Kim, D. et al. DNA skybridge: 3D structure producing a light sheet for high-throughput single-molecule imaging. Nucleic Acids Res. 47, e107 (2019).

62. Prindle, M. J. & Loeb, L. A. DNA polymerase delta in DNA replication and genome maintenance. Env. Mol Mutagen 53, 666–682 (2012).

63. Aleksandrov, R. et al. Protein Dynamics in Complex DNA Lesions. Mol. Cell 69, 1046–1061.e5 (2018).

64. Trakselis, M. A., Cranford, M. T. & Chu, A. M. Coordination and Substitution of DNA Polymerases in Response to Genomic Obstacles. Chem. Res. Toxicol. 30, 1956–1971 (2017).

65. Zhou, Y., Meng, X., Zhang, S., Lee, E. Y. C. & Lee, M. Y. W. T. Characterization of Human DNA Polymerase Delta and Its Subassemblies Reconstituted by Expression in the Multibac System. PLoS One 7, e39156 (2012).

66. Hishiki, A. et al. Structural basis for novel interactions between human translesion synthesis polymerases and proliferating cell nuclear antigen. J. Biol. Chem. 284, 10552–10560 (2009).

67. Indiani, C., McInerney, P., Georgescu, R., Goodman, M. F. & O’Donnell, M. A sliding-clamp toolbelt binds high- and low-fidelity DNA polymerases simultaneously. Mol. Cell 19, 805–815 (2005).

68. Kath, J. E. et al. Exchange between Escherichia coli polymerases II and III on a processivity clamp. Nucleic Acids Res. 44, 1681–1690 (2015).

69. Cranford, M. T., Chu, A. M., Baguley, J. K., Bauer, R. J. & Trakselis, M. A. Characterization of a coupled DNA replication and translesion synthesis polymerase supraholoenzyme from archaea. Nucleic Acids Res. 45, 8329–8340 (2017).

70. Tehseen, M. et al. Proliferating cell nuclear antigen-agarose column: A tag-free and tag-dependent tool for protein purification affinity chromatography. J. Chromatogr. A 1602, 341–349 (2019).

71. Zivanov, J. et al. New tools for automated high-resolution cryo-EM structure determination in RELION-3. Elife 7, e42166 (2018).

72. Rohou, A. & Grigorieff, N. CTFFIND4: Fast and accurate defocus estimation from electron micrographs. J. Struct. Biol. 192, 216–221 (2015).

73. Zheng, S. Q. et al. MotionCor2: anisotropic correction of beam-induced motion for improved cryo-electron microscopy. Nat. Methods 14, 331–332 (2017).

74. Zhang, K. Gctf: Real-time CTF determination and correction. J. Struct. Biol. 193, 1–12 (2016).

75. Pettersen, E. F. et al. UCSF Chimera — A Visualization System for Exploratory Research and Analysis. J. Comput. Chem. 25, 1605–12 (2004).

76. Emsley, P., Lohkamp, B., Scott, W. G. & Cowtan, K. Features and development of Coot. Acta Crystallogr. Sect. D Biol. Crystallogr. D66, 486–501 (2010).

77. Adams, P. D. et al. PHENIX: A comprehensive Python-based system for macromolecular structure solution. Acta Crystallogr. Sect. D Biol. Crystallogr. D66, 213–221 (2010).

78. Van Dijk, M. & Bonvin, A. M. J. J. 3D-DART: a DNA structure modelling server. Nucleic Acids Res. 37, W235–239 (2009).

79. Martí-Renom, M. A. et al. Comparative Protein Structure Modeling of Genes and Genomes. Annu. Rev. Biophys. Biomol. Struct. 29, 291–325 (2000).

80. Eswar, N. et al. Comparative protein structure modeling using Modeller. Curr. Protoc. Bioinforma. Chapter 5, Unit-5.6 (2006).

81. Piana, S., Robustelli, P., Tan, D., Chen, S. & Shaw, D. E. Development of a Force Field for the Simulation of Single-Chain Proteins and Protein-Protein Complexes. J. Chem. Theory Comput. 16, 2494–2507 (2020).

82. Lemkul, J. From Proteins to Perturbed Hamiltonians: A Suite of Tutorials for the GROMACS-2018 Molecular Simulation Package [Article v1.0]. Living J. Comput. Mol. Sci. 1, 5068 (2018).

83. Van Der Spoel, D. et al. GROMACS: fast, flexible, and free. J. Comput. Chem. 26, 1701–1718 (2005).

84. Lindahl, Abraham, Hess & Spoel, V. Der. GROMACS 2019.4 Source code. (Zenodo, 2019). doi: 10.5281/zenodo.3460414. (2019).

85. Abraham, M. J. et al. GROMACS: High performance molecular simulations through multi-level parallelism from laptops to supercomputers. SoftwareX 1–2, 19–25 (2015).

86. McGibbon, R. T. et al. MDTraj: A Modern Open Library for the Analysis of Molecular Dynamics Trajectories. Biophys. J. 109, 1528–1532 (2015).

87. Pedregosa, F., Weiss, R. & Brucher, M. Scikit-learn : Machine Learning in Python. J. Mach. Learn. Res. 12, 2825–2830 (2011).

88. Hunter, J. D. Matplotlib: A 2D Graphics Environment. Comput. Sci. Eng. 9, 90–95 (2007).

89. Tiberti, M., Papaleo, E., Bengtsen, T., Boomsma, W. & Lindorff-Larsen, K. ENCORE: Software for Quantitative Ensemble Comparison. PLoS Comput. Biol. 11, e1004415 (2015).

90. Humphrey, W., Dalke, A. & Schulten, K. VMD: visual molecular dynamics. J. Mol. Graph. 14, 27–28,33-38 (1996).

91. The PyMOL Molecular Graphics System, Version 2.0 Schrödinger, LLC.

92. Zhang, Z. et al. Structure of monoubiquitinated PCNA: Implications for DNA polymerase switching and Okazaki fragment maturation. Cell Cycle 11, 2128–2136 (2012).

93. Soding, J., Biegert, A. & Lupas, A. N. The HHpred interactive server for protein homology detection and structure prediction. Nucleic Acids Res. 33, W244–248 (2005).

