## Supplementary Information for "Cryo-EM structure of Pol κ−DNA−PCNA holoenzyme and implications for polymerase switching in DNA lesion bypass"

Includes:

Supplementary Figures 1-9

Description of Supplementary Movies 1-5

Supplementary Table 1

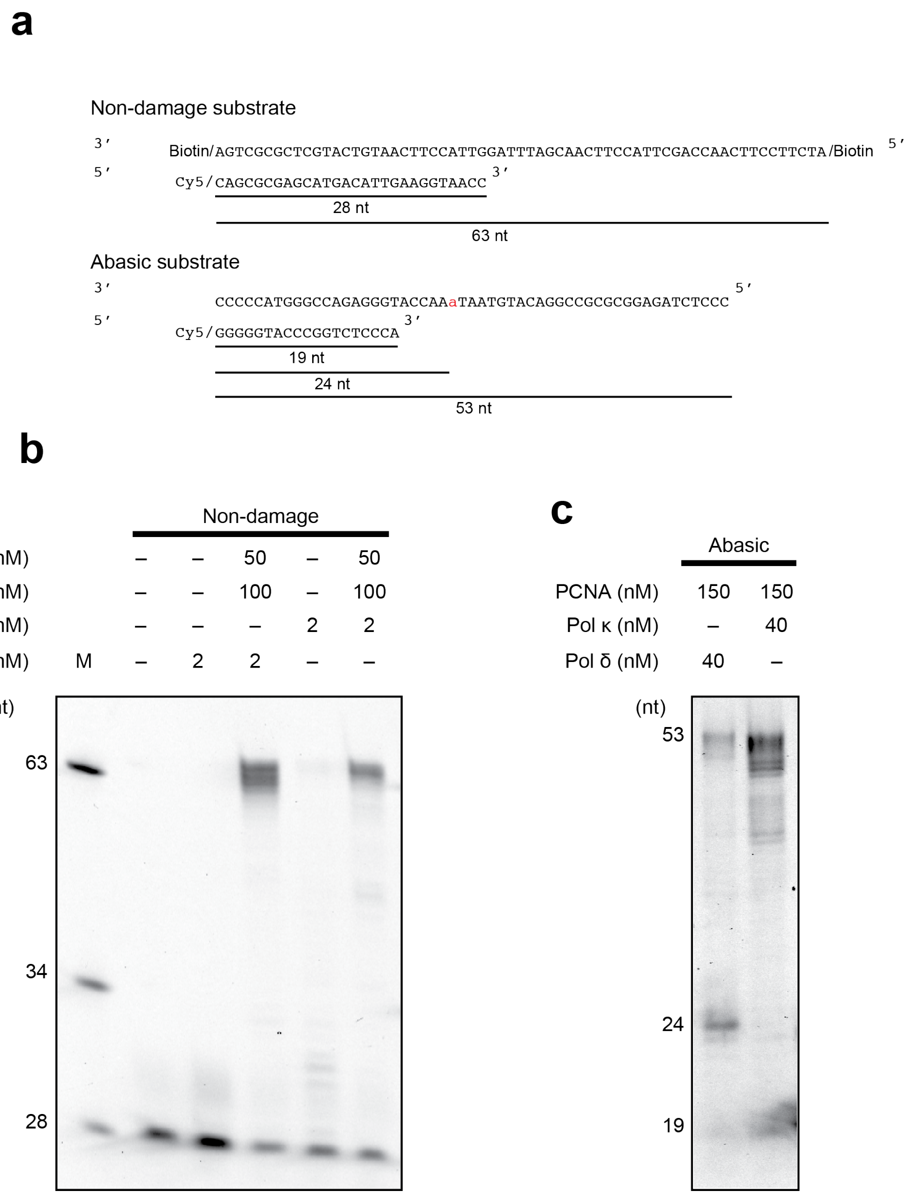


**Supplementary Figure 1.** Primer extension assay by Pol δ and Pol κ on the non-damage and abasic substrates. **a)** Schematic representation of the substrates used for the primer extension assay. **b)** Activity of Pol δ and Pol κ and their stimulation by PCNA on the non-damage substrate. Pol δ or Pol κ were incubated with the substrate in the presence of PCNA at 30 ˚C for 5 mins. **c)** Pol κ activity on the abasic substrate. Pol δ or Pol κ were incubated with the substrate in the presence of PCNA at 30 ˚C for 2 mins. The reaction's procedure is detailed in the Materials and Methods section. Protein concentrations used for the assays are indicated.

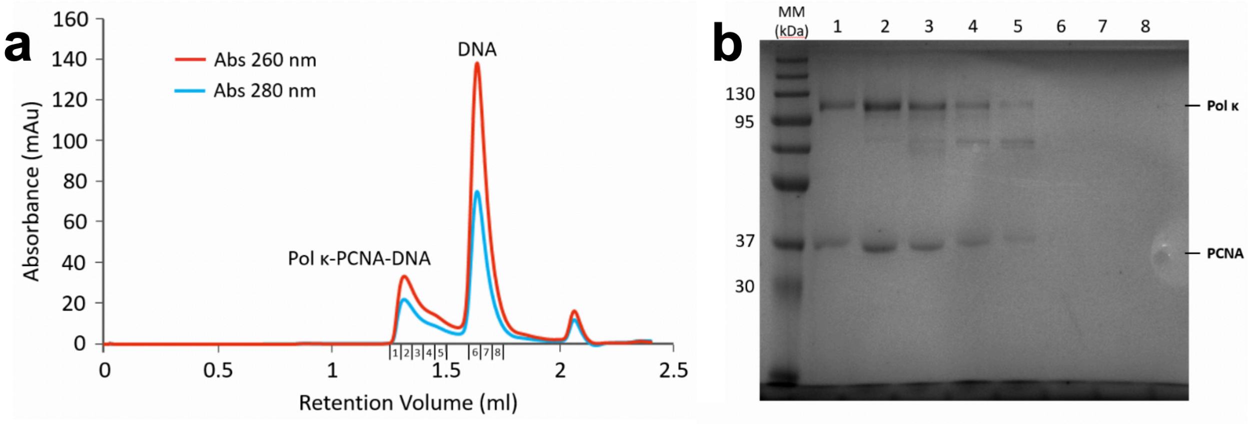


**Supplementary Figure 2**. Sample separation of the Pol κ–DNA–PCNA complex. **a)** Gel filtration chromatography of the reconstituted Pol κ–DNA–PCNA complex. **b)** The numbered peak fractions in a) were analysed by SDS-PAGE (lanes 1–8). Proteins corresponding to the bands are labeled on the right. Molecular weight standards are shown on the left.

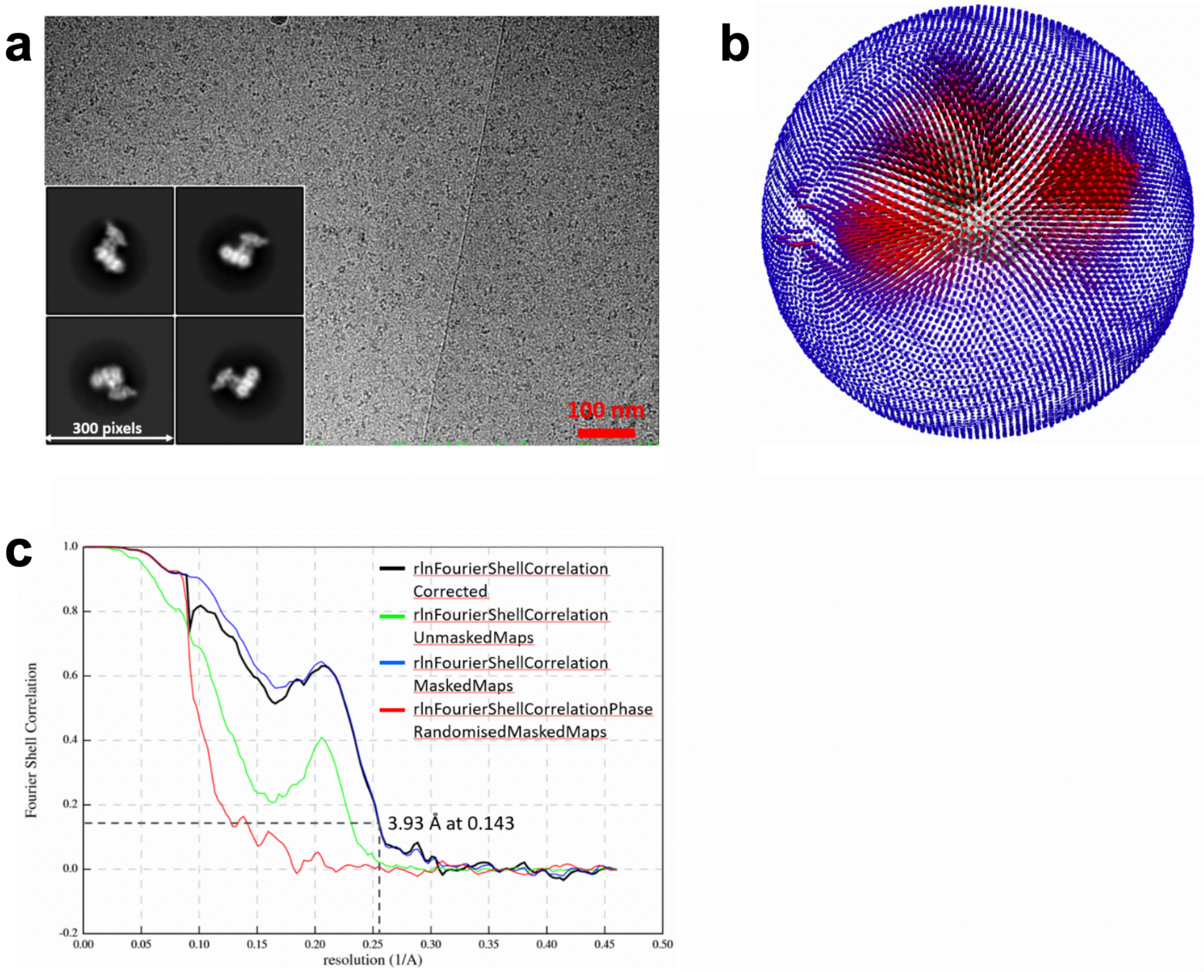


**Supplementary Figure 3.** Cryo-EM of the Pol κ holoenzyme. **a)** Electron micrograph (aligned sum) acquired using a Gatan K3 direct electron detector in super resolution mode, and representative 2D class averages. **b)** Angular distribution of projections. **c)** Gold-standard Fourier shell correlation, and resolution estimation using the 0.143 criterion.

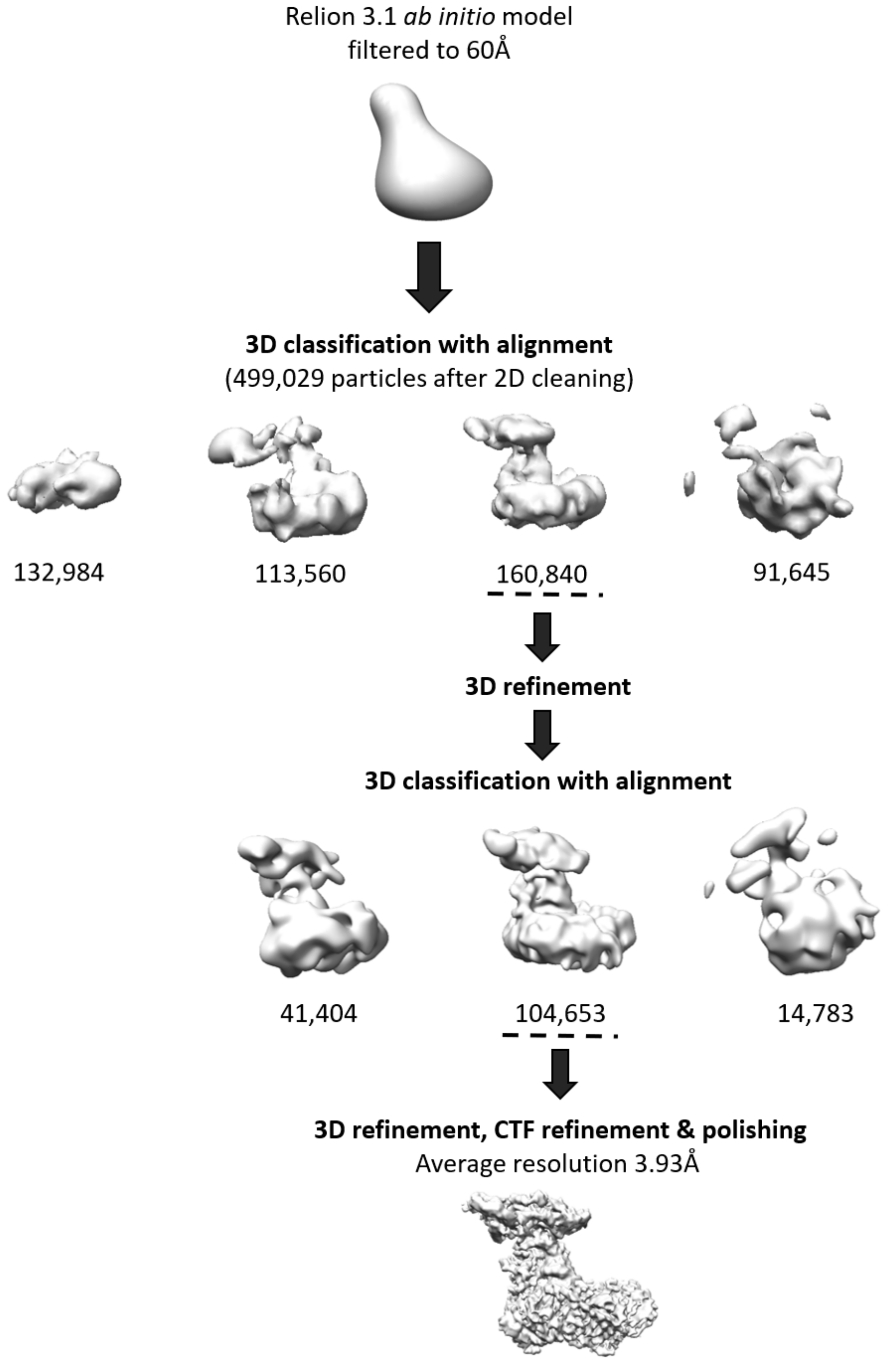


**Supplementary Figure 4.** Overview of image processing of the Pol κ holoenzyme.

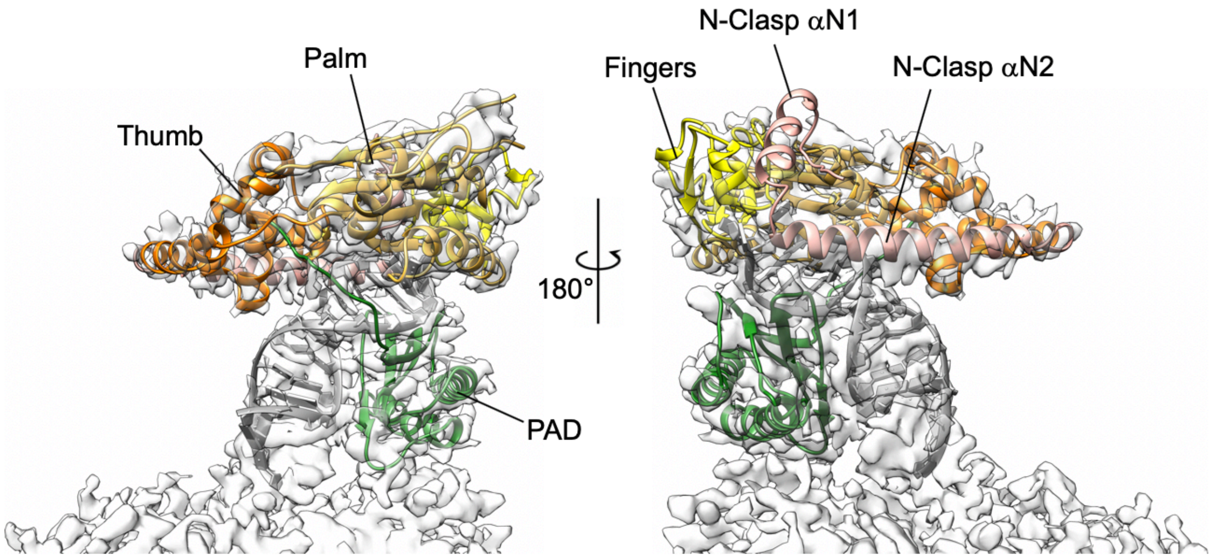


**Supplementary Figure 5.** Fitting of the X-ray structure of Pol κ ternary complex (PDB ID 2OH2) into the Cryo-EM map of Pol κ holoenzyme.

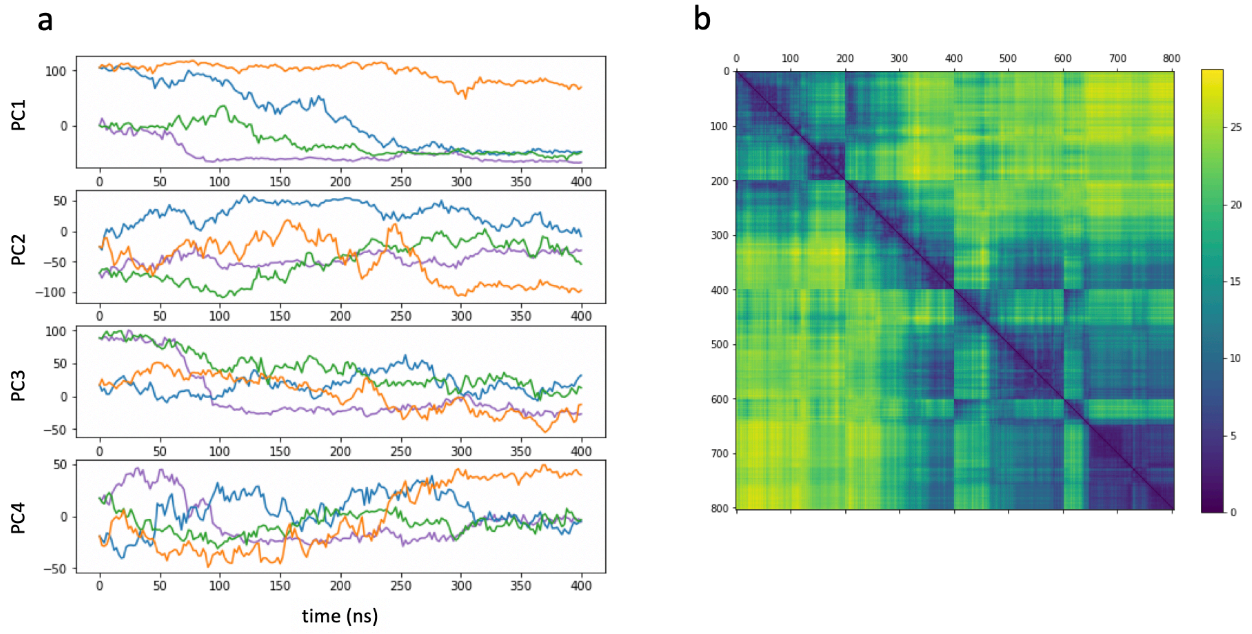


**Supplementary Figure 6. (a)** Time evolution of the projections of the four MD trajectories onto the first 4 principal components (PC). Trace color code is: Orange, apo1; Blue, apo1b; Green, apo2; Purple, apo2b. **(b)** Cross RMSD in Å of all frames in all 4 trajectories. The 800 structures represent the 200 frames of each of the 4 trajectories appended one after the other. The order of the trajectories is apo1, apo1b, apo2, apo2b. The figure shows that, although no trajectory samples the full configurational space, because trajectories mainly resemble to themselves, some trajectories sample conformations similar to others. In particular, apo1b (structures from 200 to 400) samples conformations that are closer to the ones sampled by trajectories starting from apo2 (structures from 400 to 800). This is seen from the low RMSD values between structures in the range of 300-400 with structures in the range from 400-800.

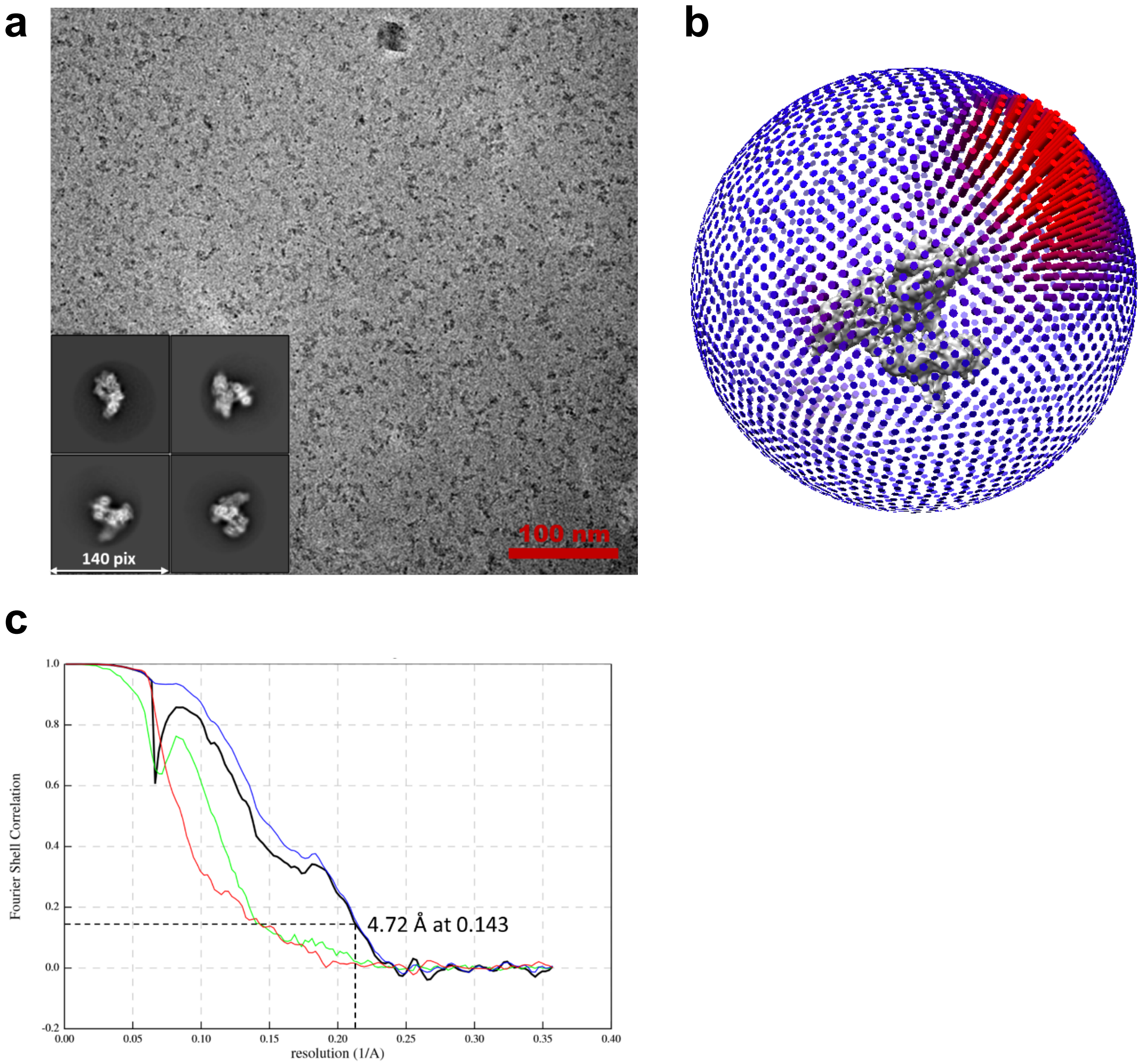


**Supplementary Figure 7**. Cryo-EM of the stalled Pol δ holoenzyme. a) Electron micrograph (aligned sum) acquired using a Gatan K2 direct electron detector in counting mode, and representative 2D class averages. b) Angular distribution of projections. c) Gold-standard Fourier shell correlation, and resolution estimation using the 0.143 criterion. Curve color code is as in Supplementary Figure 2.

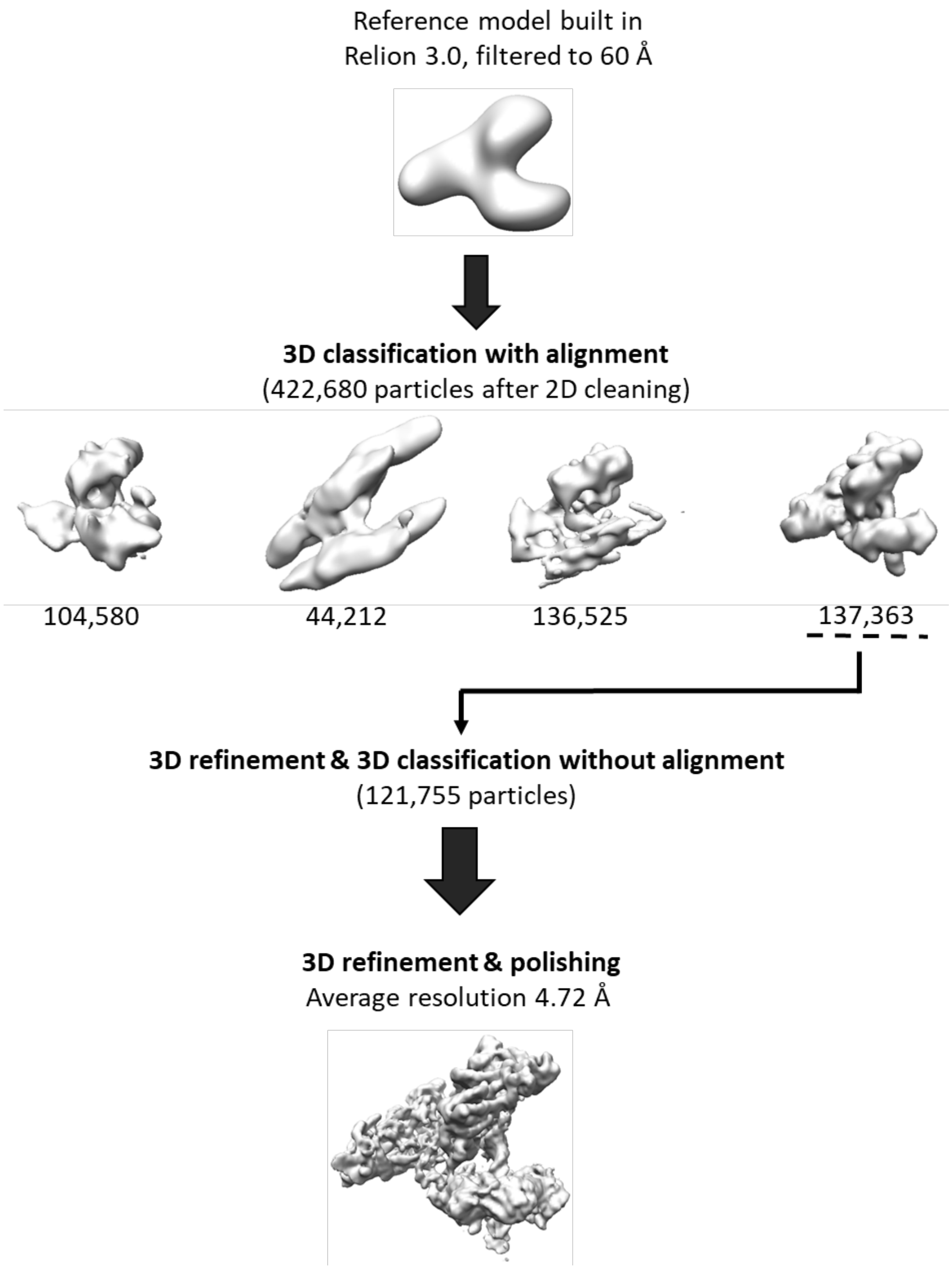


**Supplementary Figure 8.** Overview of image processing of the stalled Pol δ holoenzyme.

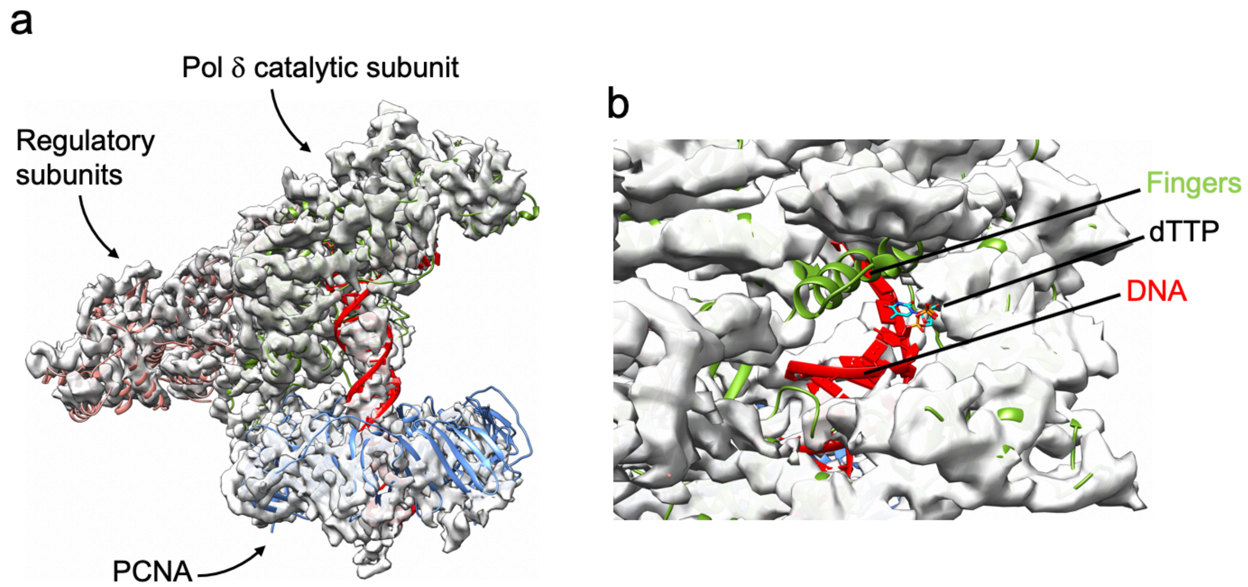


**Supplementary Figure 9. a)** Fitting of model of replicating Pol δ holoenzyme in ribbon representation (PDB ID 6TNY) into the Cryo-EM map of the stalled Pol δ holoenzyme. The fitting was performed as follows: model of stalled Pol δ was fitted into the Cryo-EM map; the model of replicating Pol δ was then aligned to that of stalled Pol δ on the catalytic domain, and the model of stalled Pol δ deleted. Poor fitting of DNA (red), PCNA (blue) and Pol δ regulatory subunits (pink) denotes conformational changes of those components in the stalled Pol δ complex. **b)** close up of the catalytic site showing the absence of density for P/T DNA and the incoming nucleotide (dTTP), and the conformational change in the fingers subdomain.

**Supplementary Movie 1.**
Apo1 MD trajectory of Pol κ–PCNA complex. Colour code: Pol κ core in bright orange; Pol κ PAD in green; PAD C-terminus in red; PCNA in skyblue.
**Supplementary Movie 2.**
Apo1b MD trajectory of Pol κ–PCNA complex. Colour code: Pol κ core in bright orange; Pol κ PAD in green; PAD C-terminus in red; PCNA in skyblue.
**Supplementary Movie 3.**
Apo2 MD trajectory of Pol κ–PCNA complex. Colour code: Pol κ core in bright orange; Pol κ PAD in green; PAD C-terminus in red; PCNA in skyblue.
**Supplementary Movie 4.**
Apo2b MD trajectory of Pol κ–PCNA complex. Colour code: Pol κ core in bright orange; Pol κ PAD in green; PAD C-terminus in red; PCNA in skyblue.
**Supplementary Movie 5.**
Morphing between replicating and stalled Pol δ–DNA–PCNA complex structures, showing the release of P/T DNA from the active site and conformational changes in Pol δ and PCNA. Color code: Pol δ p125 core in forest green, with the thumb domain in bright green;

Pol δ p125 CTD in orange; p50 subunit in pink; p66 subunit in yellow; p12 subunit in red; PCNA in blue.

**Supplementary Table 1.** Cryo-EM data collection, refinement and validation statistics

| | Polκ-DNA-dTTP-PCNA (EMD-11291) (PDB 6ZMH) | Polδ-DNA-PCNA (EMD-11290) (PDB 6ZMF) |
|---|---|---|
| **Data collection and processing** | | |
| Magnification | 81,000 | 105,000 |
| Voltage (kV) | 300 | 300 |
| Electron exposure (e–/Å$^2$) | 47 | 48 |
| Defocus range (μm) | -0.7 to -1.5 | -0.6 |
| Pixel size (Å) | 1.086 | 1.4 |
| Symmetry imposed | C1 | C1 |
| Initial particle images (no.) | 2,285,897 | 3,189,106 |
| Final particle images (no.) | 104,653 | 121,755 |
| Map resolution (Å) | 3.93 | 4.72 |
| FSC threshold | 0.143 | 0.143 |
| Map resolution range (Å) | 3.68-5.86 | 4.32-13.90 |
| **Refinement** | | |
| Initial model used (PDB code) | 2OH2, 6TNY | 6TNY |
| Model resolution (Å) | 3.9 | 4.7 |
| FSC threshold | 0.143 | 0.143 |
| Model resolution range (Å) | N/A | N/A |
| Map sharpening *B* factor (Å$^2$) | -93.5 | -145.6 |
| Model composition | | |
| Non-hydrogen atoms | 10112 | 19746 |
| Protein residues | 1180 | 2393 |
| Nucleotide residues | 53 | 54 |
| Ligands | 1 | 2 |
| R.m.s. deviations | | |
| Bond lengths (Å) | 0.008 | 0.007 |
| Bond angles (°) | 0.965 | 1.156 |
| Validation | | |
| MolProbity score | 1.72 | 2.09 |
| Clashscore | 6.12 | 12.36 |
| Poor rotamers (%) | 0.51 | 0.47 |
| Ramachandran plot | | |
| Favored (%) | 94.3 | 92.92 |
| Allowed (%) | 5.70 | 6.61 |
| Disallowed (%) | 0.00 | 1.12 |
| Model vs Data | | |
| CC (mask) | 0.69 | 0.58 |
| CC (box) | 0.74 | 0.73 |
| CC (peaks) | 0.60 | 0.53 |
| CC (volume) | 0.69 | 0.60 |
